# Architecture of the Serotonergic Projectome in the Mouse Brain

**DOI:** 10.64898/2026.07.15.738594

**Authors:** Jun Ho Song, Drew Friedmann, Yunming Wu, Ashley Moses, Jalal Kenji Baruni, Yuan Yuan, Isabella Rana, Kelly Zhao, Tom Hindmarsh Sten, Qian Wang, Shuyun Alina Xiao, Xiaoke Chen, Scott W. Linderman, Liqun Luo

## Abstract

The serotonin system innervates nearly the entire brain to support diverse functions, and its dysfunction is implicated in multiple psychiatric disorders. Although recent studies suggested that this anatomically diffuse system supports differentiated rather than uniform modulation, its overall organization is unclear. Here, using systematic whole-brain axon tracing and integrated analyses, we show that the serotonin projectome is organized by functional relatedness rather than physical proximity of its targets. Dorsal and median raphe serotonin neurons partition into five projectomic groups, each preferentially innervating the hippocampus, basal ganglia, cortex, medial interbrain, or brainstem/lateral thalamus, with within-group structure ranging from discrete subgroups to continuous variation. Spatial transcriptomic analyses reveal that each projectomic group corresponds to a distinct combination of transcriptomic clusters, with transcriptome and somatic position jointly predicting projectomic identity. Projection-specific serotonin depletion produces dissociable behavioral phenotypes. Together, projectomic identity emerges as a central axis linking axon collateralization, molecular diversity, and behavioral function.

## INTRODUCTION

Serotonin (5-hydroxytryptamine, 5HT) is an ancient neuromodulator widely employed across the animal kingdom. It plays diverse roles across species, from mediating sensitization of the gill-withdrawal reflex in *Aplysia*^1^ and regulating foraging in *C. elegans*^2^ to nutrient intake regulation in *Drosophila*^3^ and locomotor learning in zebrafish^4^. In mammals, serotonin has been implicated in diverse physiological, behavioral, and cognitive functions^5^, including reward^6,7^, patience^8,9^, learning^10^, seasonal circadian rhythm^11^, anxiety-like behaviors^12,13^, and social behaviors^14,15^. The clinical importance of serotonergic signaling is underscored by the widespread use of SSRIs to treat psychiatric disorders including depression, anxiety, and obsessive-compulsive disorder^16–18^.

In the mammalian brain, serotonin neurons are clustered within a few discrete nuclei in the brainstem, mainly the dorsal and median raphe nuclei (DR and MR)^19^. Despite comprising a minuscule fraction of the neuronal population (about 1 serotonin neuron for every 10^4^ or 10^6^ neurons in the mouse or human brain, respectively^20,21^), axons of serotonin neurons arborize across nearly the entire brain^19,22,23^. While most experiments exploring the physiological and behavioral functions of serotonin neurons have been conducted by treating serotonin neurons (mostly from DR) as a monolith, recent work has underscored that the raphe nuclei contain transcriptomically and anatomically diverse populations of serotonin neurons that stem from distinct developmental lineages and exhibit unique functional properties^24–28^. For example, DR serotonin neurons that project to the orbitofrontal cortex and central amygdala have distinct anatomical, physiological, and functional characteristics^24^. However, these two subsystems represent only a small fraction of raphe serotonin neurons.

Given that projection targets are central to the physiological and behavioral functions of serotonin neurons, these studies raised fundamental questions about the organization of the serotonergic projectome. How many distinct serotonin subsystems constitute the entire serotonin projectome in the mouse brain? Which target regions are co-innervated by the same serotonin neurons, and which are innervated by distinct ones? How does the transcriptomic diversity of serotonin neurons^25–27^ map onto this projectomic structure? How do distinct projectomic populations contribute to behavior? Here, we combine viral-genetic tracing, whole-brain imaging, spatial transcriptomics, projection-specific loss of serotonin function, and an integrated analytic framework to address these questions.

## RESULTS AND DISCUSSION

### Determining whole-brain serotonin axon collateralization patterns

To investigate the extent to which serotonin neurons can be partitioned based on their whole-brain projection patterns, we implemented an intersectional viral-genetic strategy to label serotonergic populations defined by individual projection targets and map their axonal collateralization. We used transgenic mice carrying a Flp and Cre double-gated tdTomato reporter (*Ai65*)^29^ and either *Sert-Cre*^30^ or *Tph2-iCreER* (**Figure 1A**; **Methods**). Into different projection targets of serotonin neurons in these mice, we injected a retrogradely transducing AAV (AAV_retro_) encoding Cre-dependent Flp recombinase^24,31^ (**Table S1**). This approach is independent of cell body position within the raphe nuclei, avoiding biases that can arise from raphe-targeted injections. Following reporter expression, we cleared^32^ and imaged the brains using light-sheet microscopy, segmented labeled axons using TrailMap^33^, and registered the data to Allen CCFv3^34^ (**Figure 1B**). This experimental pipeline enabled us to directly compare whole-brain serotonergic axon projection data obtained from different mice in the same reference space.

**Figure 1.**
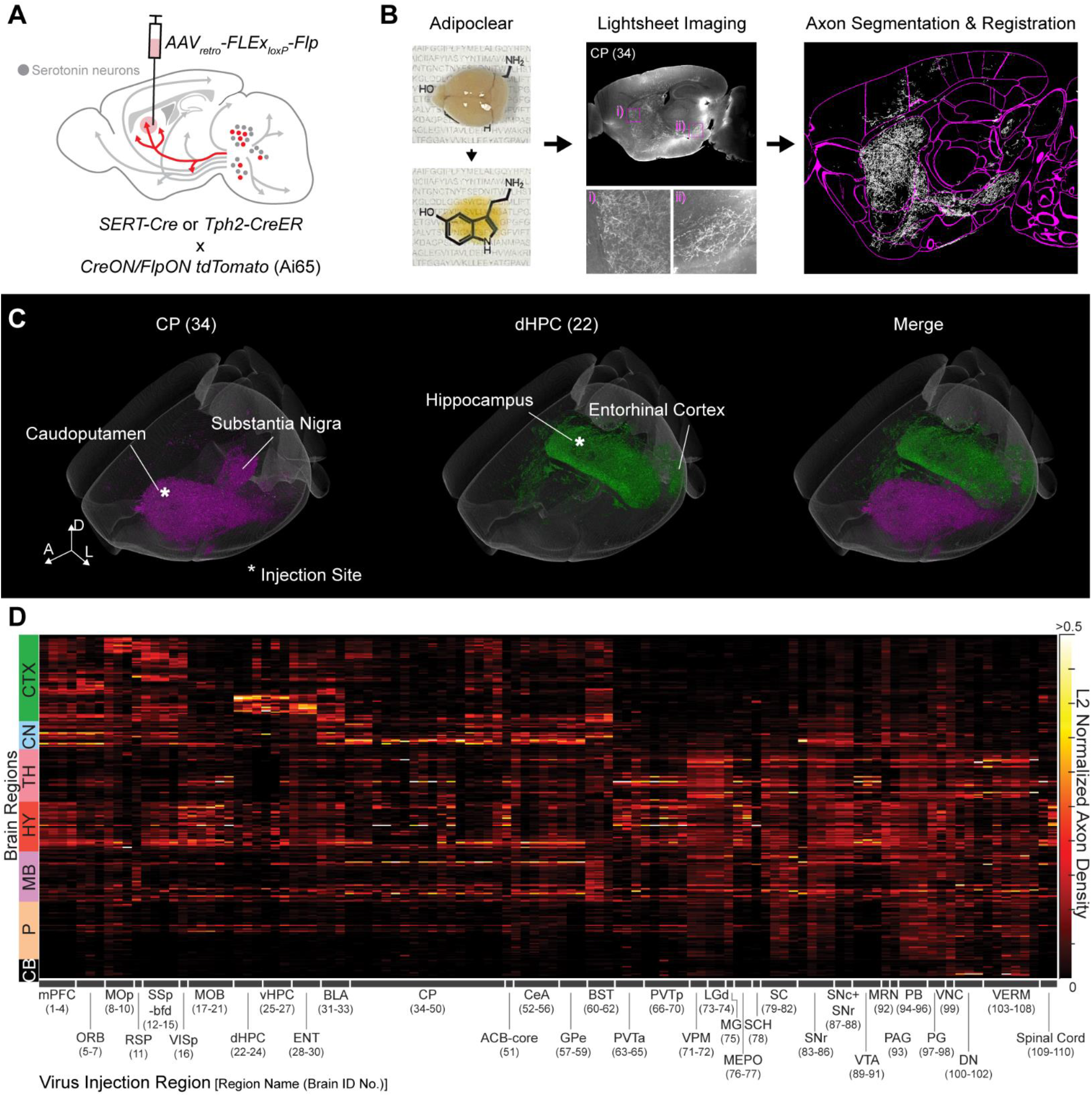
Whole-brain axon collateralization of serotonin neurons in DR and MR. (**A**) Schematic of retrograde labelling of projection-specific serotonin neurons. (**B**) Brain sample processing pipeline. The left panels show a perfused brain (top) becoming transparent (bottom) after Adipoclear^32^ and tdTomato staining. The middle panels show a raw image from light-sheet imaging of a sagittally oriented brain (top) and a zoomed-in view of regions containing labeled axons (bottom). CP denotes the virus injection site (brain ID in parenthesis). Right, voxels containing labeled axons (middle) were segmented and registered to Allen CCFv3. White, segmented axons; magenta lines, brain region boundaries. (**C**) 3-D views of segmented and registered axons from two example brains and their merged representation. Axis labels: A, anterior; D, dorsal; L, lateral. *, AAV_retro_ injection sites. (**D**) L2-normalized axon density in 280 brain regions. Brains (columns, ID in parenthesis) injected at various regions (*x*-axis; coordinates in **Table S1**) ordered from left to right according to the structural hierarchy of CCFv3. Full names of brain regions in **Table S2**. Axon densities in 280 brain regions (rows) were quantified. See **Figure S1**, **Tables S1**, and **S2** for additional data.

**Figure 1C** and **Video S1** exemplify contrasting whole-brain collateralization patterns of serotonin neurons when we initiated labeling (via AAV_retro_ injection) at two sites—the dorsomedial caudoputamen (CP) or the dorsal hippocampus (dHPC). The injection into the CP densely labeled basal ganglia nuclei, including the CP, globus pallidus, and substantia nigra reticulata (SNr), whereas the injection into the dHPC instead labeled the broader hippocampal network, including ventral hippocampus (vHPC) and entorhinal cortex (ENT), with minimal overlap between the two examples.

To globally assess the collateralization patterns of serotonin neurons, we collected imaging data from 110 brains that were injected into 35 distinct, well-studied regions with minimal virus spillover into neighboring regions (**Figure S1A**). Collectively, the labeled axons across these brains covered all brain regions known to be innervated by serotonin neurons^23^ (**Figure S1B–K**). We then quantified the density and volume of labeled axon in 280 divisions of the entire mouse brain using the Allen CCFv3 segmentation^34^ (**Figure 1D; Table S2**). Injections at the same brain region produced similar collateralization patterns, whereas injections at different brain regions yielded varying degrees of similarity depending on the injection sites. This suggests that the serotonin projectome could potentially be deconstructed into distinct groups based on their shared target regions.

### DR and MR serotonin neurons comprise five projectomic groups

Although **Figure 1D** provides a rich dataset on the architecture of the serotonin projectome, examining shared organizational features across mice is challenging in this high-dimensional space (110 mice × 280 regions). To extract shared projection patterns (henceforth “basis patterns”, analogous to principal components in principal component analysis), we applied non-negative matrix factorization (NMF)—a technique widely used to extract common patterns in large datasets with non-negative values^35–39^—followed by clustering of individual brains in the reduced space (**Figure 2A, B**; **Methods**). To determine the dimensionality of the latent space—the optimal number of basis patterns *k*—we performed random “speckled” cross-validation with varying values of *k* (**Figure 2C**). Reconstruction error on held-out data (RMSE_Test_) decreased with increasing *k* until plateauing at *k =* 5, where it reached the best-achievable baseline (RMSE_Replicate_; defined by within-site animal-to-animal variability) and remained well below chance (RMSE_Shuffle_). This suggests that a low-dimensional representation with five basis patterns effectively captures the dominant structure in the dataset.

**Figure 2.**
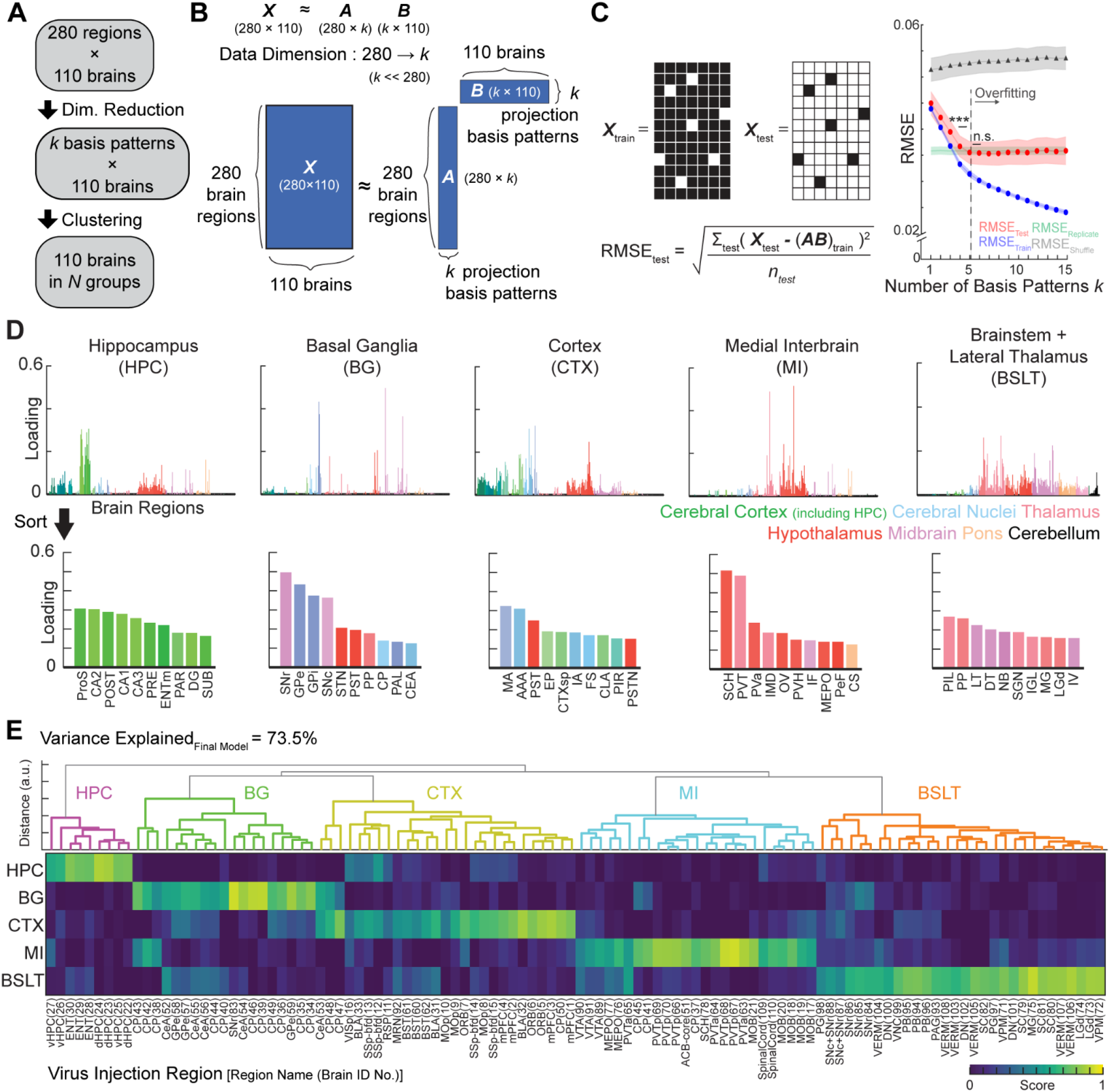
Five projectomic groups of serotonin neurons identified through NMF analysis. **(A)** Analysis pipeline. The matrix in Figure 1D (**Table S2**) is first reduced to *k* dimensions to enhance feature extraction and reduce noise. The samples in the *k*-dimensional space are then clustered into *N* projectomic groups. **(B)** Schematic of dimensionality reduction using non-negative matrix factorization (NMF). ***X***, matrix containing the normalized axon density data in Figure 1D; ***A***, projection basis matrix; ***B***, score matrix (i.e., the dimensionality-reduced dataset). **(C)** Left, 10% of the elements in ***X*** were randomly assigned to ***X_test_***, and the remainder to ***X_train_***. Right, for each *k*, reconstruction error from “speckled” cross-validation was calculated using root-mean-square error (RMSE) (1,000 iterations). RMSE_Replicate_ denotes the best-achievable performance based on inter-replicate variability. RMSE_Shuffle_ denotes the baseline performance expected by chance. One-way ANOVA followed by Bonferroni correction. Error bars, standard deviation. ***, *p* < 0.001; n.s., not statistically significant. **(D)** Top, Loadings in the five projection basis patterns across the 280 brain regions (i.e., matrix ***A*** in panel A). Bottom, top 10 brain regions per basis pattern (full names in **Table S2**) sorted by loading. **(E)** Re-description of axon density data (Figure 1D) with the five basis patterns. The 110 brains were hierarchically clustered according to their scores in the dimensionality-reduced data points (i.e., matrix ***B*** in panel A). A score for a particular basis pattern represents the weight a basis pattern contributes to reconstructing a data point in its original 280-D space. Sample names follow the notation used in Figure 1D. The optimal cluster number determined by gap statistic (**Figure S2B**). a.u., arbitrary unit. See **Figure S2–S4** and **Table S2** for additional data.

Each of the resulting five basis patterns (matrix ***A*** in **Figure 2B**) captured a distinct anatomical pattern of projections (**Figure 2D**): hippocampal-entorhinal network (basis 1), basal ganglia (basis 2), cortical regions (basis 3), medial thalamus and hypothalamus (collectively, medial interbrain; basis 4), and brainstem and lateral thalamic nuclei including sensory thalamus such as the dorsal lateral geniculate (LGd) and medial geniculate body (MG) (basis 5). Based on the anatomical characteristics of these basis patterns, we hereafter refer to them as the hippocampal (HPC), basal ganglia (BG), cortical (CTX), medial interbrain (MI), and the brainstem-and-lateral-thalamic (BSLT) patterns.

Having identified a low-dimensional representation of serotonergic projections, we hierarchically clustered individual brains using their basis scores (matrix ***B*** in **Figure 2B**), yielding five projectomic groups (**Figure 2E**). Each group was characterized by a dominant score on one of the five basis patterns, and we named the groups accordingly (HPC, BG, CTX, MI, BSLT). We note that the number of projectomic groups (*N*) need not equal the number of bases (*k*); a group could exhibit high scores on multiple bases, and multiple groups could share a dominant basis. Our data, however, were best described by the simplest mapping of one group per basis.

The dimensionality-reduced dataset reflected each brain’s serotonergic projection characteristics. For example, the CP-injected (ID 34) or the dHPC-injected (ID 22) brain in **Figure 1C** had a dominant BG or HPC score and was thus classified into the BG or HPC group, respectively. A subset of BG-injected brains, however, showed high scores on both BG and CTX bases (e.g., IDs 48, 47, 50; **Figure 2E**), suggesting that CP subregions may be co-innervated by distinct projectomic populations—a point we will return to later.

To further validate this five-group framework, we first confirmed its robustness to analytical choices: similar groupings were obtained using the original 280-D data (**Figure S2A**), alternative clustering methods (**Figure S2B, C**), subsampled data (**Figure S2D**), and axon volume instead of density (**Figure S3**; **Table S2**). We next tested whether brains within the same group share more similar collateralization patterns than brains from different groups. Pairwise cosine similarity between the 110 brains (based on 280-D projection density; **Figure 1D**) revealed five high-similarity squares along the diagonal, closely mirroring our projectomic grouping (**Figure S4A**). We further compared brains with injection sites at (i) the same regions, (ii) different regions within the same projectomic group, and (iii) different regions from other projectomic groups (**Figure S4B**). We found that projection similarity (ii) did not significantly differ from same-region similarity (i) for all groups except CTX and significantly exceeded between-group similarity (iii) across all groups, demonstrating that the projectomic groups represent coherent, broadly collateralized projection systems.

Interestingly, the cosine similarity in (iii) was substantially above zero. This could be attributable to: (1) different projectomic groups share some target regions (**Figure 2D**), and (2) some individual brains contributed to multiple bases (**Figure 2E**). This observation parallels gene expression across cell types: individual brain regions can be innervated by multiple projectomic groups, but a subset that receives dominant input from a single group can serve as “marker regions”, analogous to marker genes (**Figure S4C, D**). These marker brain regions provide anatomical access to specific projectomic groups for functional studies, as we demonstrate later. Together, the five projectomic groups form a hybrid organization—discrete group-level structure alongside partial target sharing—rather than tiling the brain in a mutually exclusive manner.

### Within-group organization: from discrete subgroups to continuous gradients

The five projectomic groups provide a first-order decomposition of the serotonin system, but within-group heterogeneity could further refine this organization. For example, the distribution of CP-injected samples across multiple projectomic groups (BG, CTX, MI; **Figure 2E**) suggested subpatterns within CP not captured by the whole-brain-level categorization. Given the broad spatial coverage of our CP injection sites (**Table S1**), we applied NMF to serotonergic axon distributions within the isolated CP volume to identify subpatterns (**Figure 3A**). Speckled cross-validation identified *k* = 4 as the optimal number of basis-subpatterns (**Figure 3B**). These subpatterns were enriched in specific CP subregions across the 3D volume (**Figure 3C**). To understand how these CP subpatterns relate to brain-wide projection differences, we correlated each CP subpattern’s score with projection to non-BG targets (**Figure S5A–C**). This revealed, for example, that cortical labeling in CP-injected brains was most strongly associated with serotonergic axons targeting ventrolateral CP.

**Figure 3.**
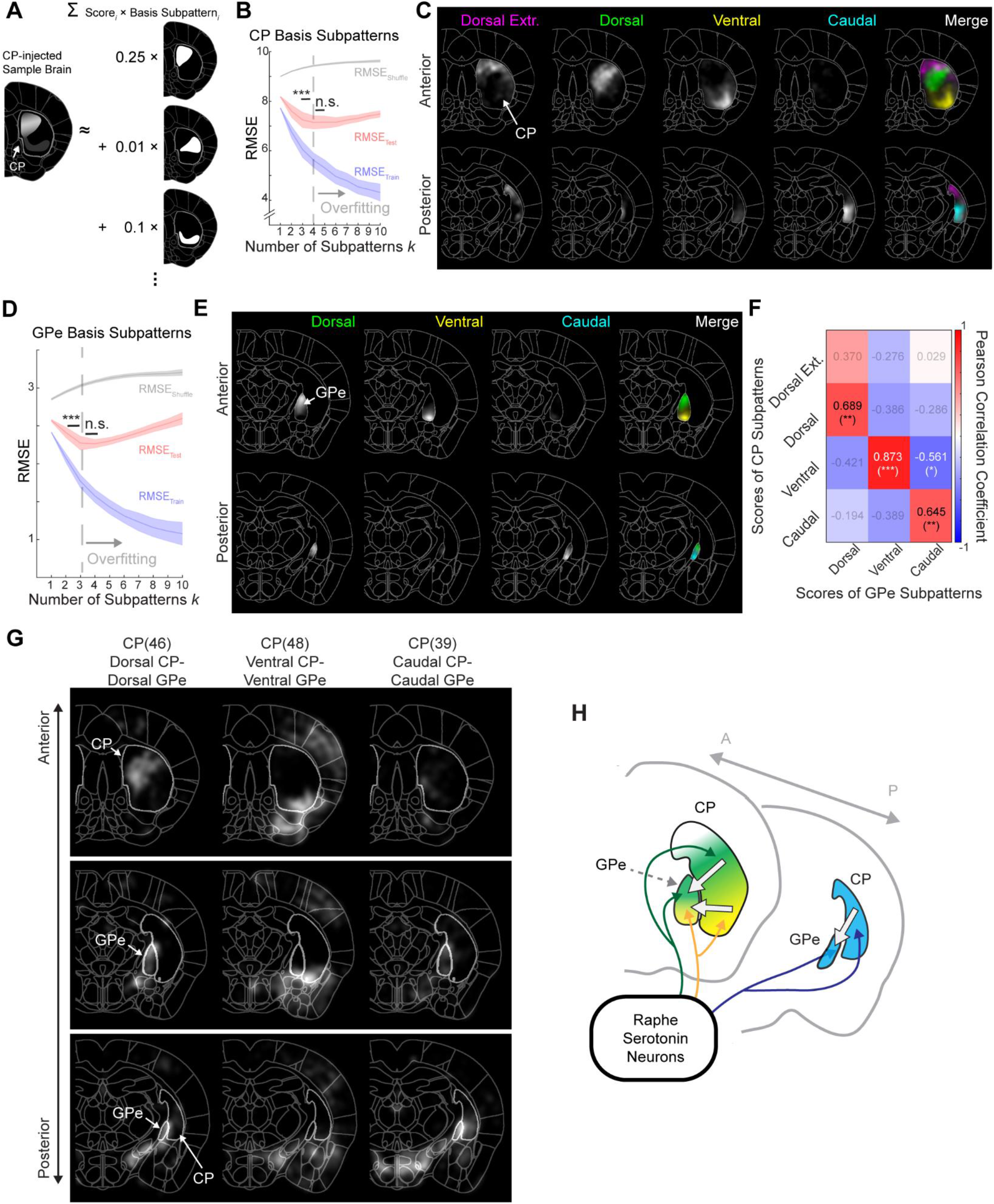
Topographic organization of serotonin projectomic subpatterns in the basal ganglia. **(A)** Analysis schematic: serotonergic CP innervation approximated as a linear combination of basis subpatterns and scores. **(B)** Cross-validation to determine k for NMF on CP volumes (n = 17 CP-injected brains). 500 iterations. One-way ANOVA with Bonferroni correction. Error bars, SD. ***p < 0.001; n.s., not significant. **(C)** Coronal views of the four extracted basis subpatterns of CP. Axon innervation patterns in the CP across the 17 brains can be reconstructed using a linear combination of these four basis patterns and the corresponding scores shown in **Figure S5A**. Subpattern names color-coded to match merge panel. **(D)** Cross-validation for NMF on GPe volumes (same brains). ***p < 0.001; n.s., not significant. **(E)** Coronal views of the three GPe basis subpatterns; scores in **Figure S5D**. **(F)** Pairwise Pearson correlation coefficient between the scores of CP- and GPe-subpatterns shown in **Figure S5A** and **S5D**. *, *p* < 0.05; **, *p* < 0.01; ***, *p* < 0.001. Values in gray indicate no statistical significance. **(G)** Coronal sections from three sample brains illustrating the topographic serotonergic innervation of BG subpatterns in CP and GPe. The brains in the left, middle, and right columns exemplify the topographic organization of serotonergic projections to dorsal CP/dorsal GPe, ventral CP/ventral GPe, and caudal CP/caudal GPe. White boundaries, CP and GPe. **(H)** Model illustrating that subpopulations of serotonin neurons innervate connected subregions of the CP and GPe. White arrows from CP to GPe represent known connectivity between these two regions.See **Figure S5** and **S6** for additional data.

The CP projects topographically to globus pallidus externus (GPe), a downstream basal ganglia region^40,41^. NMF analysis revealed that the GPe was optimally segregated into three subpatterns (**Figure 3D, E**). Correlating CP and GPe subpattern scores (**Figure S5A, D**) revealed topographically arranged serotonergic co-innervation: dorsomedial CP with dorsal GPe, ventral CP with ventral GPe, and caudal CP with caudal GPe (**Figure 3F, G**). This mirrors the known CP→GPe topography^40^, suggesting that parallel CP→GPe subcircuits within the basal ganglia are each preferentially co-innervated by separable subpopulations of serotonin neurons within the BG projectomic group (**Figure 3H**).

We next asked whether the CTX group exhibits similar substructure. The 27 CTX-group brains showed substantial variability in isocortical axon density (**Figure S5E**), consistent with the notable same-region vs. within-group similarity gap observed in this group (**Figure S4B**). Applying NMF to isocortical axon density across these 27 brains identified *k* = 3 as optimal (**Figure S5F, G**). Unlike the CP-injected brains, CTX-group brains did not form distinct clusters in the 3D space, and instead varied continuously (**Figure S5H, I**). Pairwise cosine similarity of neocortical axon density was weakly but significantly anti-correlated with physical distance (**Figure S5J**), suggesting that an adjacency effect explains a small fraction of the variation.

NMF analysis of the remaining three groups did not reveal statistically significant subpatterns for MI or BSLT (**Figure S6A**). The HPC pattern showed modest improvement at *k* = 2, splitting into two subpatterns projecting primarily to hippocampal areas and retrohippocampal regions (entorhinal cortex), respectively (**Figure S6B**, **C**). Together, these analyses reveal a dual organization of serotonergic innervation, from discrete subgroups in the basal ganglia to continuously varying projections across the cortex.

### Organization of the mouse brain from the perspective of serotonin modulation

Our data thus far suggest that one organizing principle of the serotonin projectome is functional relatedness rather than physical proximity. This is exemplified by the HPC group, which predominantly innervates the hippocampal subdivisions (**Figure 2D**)—structures that play crucial roles in cognitive representations of space, time, and memories^42–44^. Its preferential association with the entorhinal cortex, but not other equally proximal cortices, further reinforces the idea of functional rather than spatial groupings. Similarly, the BG group innervates the anatomically distributed but functionally integrated dorsal striatal pathway (CP, GPe, GPi, STN, SNr)^45,46^. The BG subgroups we identified exhibit further refinement in their preferential innervation of parallel CP→GPe subcircuits (**Figure 3**), reinforcing functional similarity as a key organizing feature.

Beyond these organizational observations, our projectome revealed several unexpected groupings—the separation of related brain regions into distinct projectomic groups and clustering of seemingly less-related brain regions into the same group (**Figure 2E**). First, central amygdala (CeA) and basolateral amygdala (BLA) belonged to distinct projectomic groups: CeA-injected brains clustered into the BG group whereas BLA-injected ones clustered into the CTX group. Although surprising given the physical adjacency and functional overlap (e.g., in regulating fear learning and emotion^47^), this is consistent with their distinct developmental origins and cellular compositions^48,49^. Second, ventral tegmental area (VTA)- and nucleus accumbens core (ACB-core)-injected brains clustered into the MI rather than the BG group. This was unexpected given the close relationship and circuit architecture of ACB (part of the ventral striatum) and CP (dorsal striatum), as well as the adjacency of VTA and SNc dopamine neurons in their cell body positions and projection patterns in the striatum. However, this dissociation is consistent with the functional divergence of these circuits: although both are striatal, VTA– ACB primarily supports motivation and reward, whereas SNc–CP supports motor control and learning^45,46,50^. Our finding suggests that serotonergic co-innervation reflects this functional divergence rather than their shared anatomical organization. Third, brains injected in subcortical motor- and sensory-related regions—such as lateral geniculate nucleus, superior colliculus, parabrachial nucleus, vermis of the cerebellar cortex, and cerebellar dentate nucleus—were classified into the BSLT group, suggesting a role of serotonergic co-innervation in sensorimotor coordination. Lastly, several seemingly unrelated regions located near the midline of the brain were commonly targeted by the MI group of serotonin neurons (**Figure 2D, E**). These include the paraventricular thalamic nuclei, main olfactory bulb, spinal cord, VTA and ACB, as well as hypothalamic nuclei with homeostatic functions such as the suprachiasmatic nucleus (circadian rhythm^11,51^), organum vasculosum of lamina terminalis and median preoptic nucleus (osmoregulation^52,53^ and thermoregulation^54^). Whether serotonergic neurons modulate these targets in unison or through branch-specific mechanisms^11^—and what functional or developmental relationships might underlie their shared innervation—remains to be determined.

### Projection-specific cell body distributions in the raphe nuclei

Our viral-genetic strategy for determining the whole-brain collateralization patterns of serotonin neurons (**Figure 1A**) is agnostic to their cell body positions. Yet, the location of these labeled cells in the DR and MR varied substantially with injection sites. For instance, mPFC-projecting neurons were broadly distributed across both regions, whereas GPe-projecting neurons preferentially clustered in DR, and nucleus accumbens core (ACB-core)-projecting neurons preferentially clustered in MR and caudal DR (cDR) (**Figure 4A**).

**Figure 4.**
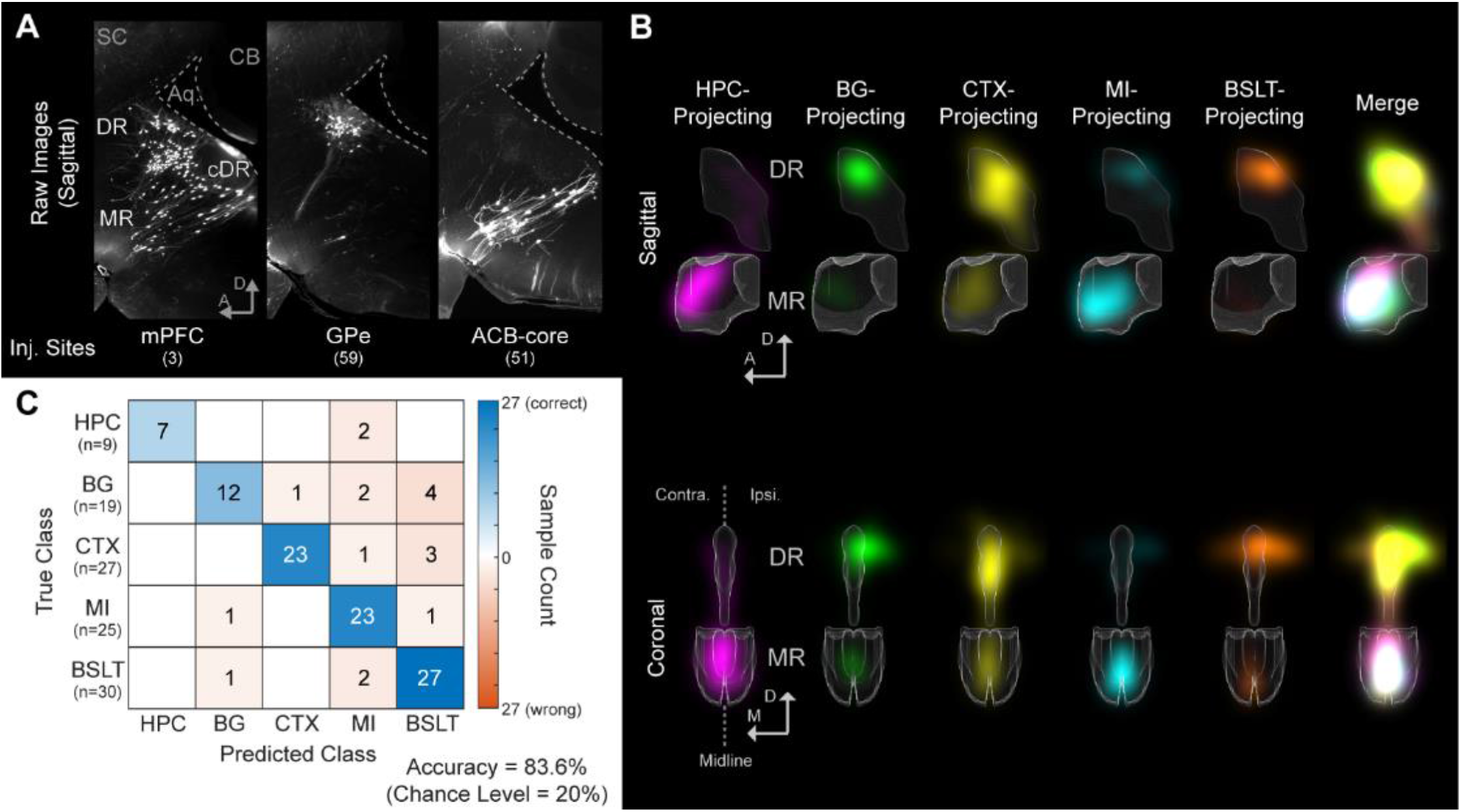
Mapping cell body positions of serotonergic neurons in the raphe nuclei by their projection groups. **(A)** Sagittal views of example brains illustrating diverse serotonergic cell body distributions in the raphe; AAV_retro_ injection sites shown at the bottom. A 100-µm-thick stack at a comparable media-lateral position was selected from each lightsheet-imaged whole-brain sample and projected by maximal intensity. Axes: A, anterior; D, dorsal. Region abbreviation: Aq, aqueduct; CB, cerebellum; cDR, caudal dorsal raphe; SC, superior colliculus. **(B)** Average distributions of projection-specific serotonergic neurons in DR and MR. Axis labels: A, anterior; D, dorsal; M, medial; Contra, contralateral; Ipsi, ipsilateral. **(C)** Confusion matrix for projectomic-group prediction from raphe cell body distributions, using a supervised NMF model and evaluated by 5-fold stratified cross-validation. Numbers, brain counts. Overall accuracy: 83.6% (theoretical chance 20%). See **Figure S7** for additional data.

Mean distributions of the five projectomic groups showed partial overlap alongside distinct anatomical biases (**Figure 4B**): HPC-projecting neurons were concentrated in anterodorsal MR with some in cDR; BG-projecting neurons clustered in ipsilateral dorsolateral DR with some in anterior MR; CTX-projecting neurons spanned the entire raphe and, uniquely, densely populated the ventral DR^24^; MI-projecting neurons were mostly in MR with some in cDR; and BSLT-projecting neurons were concentrated in ipsilateral dorsolateral DR with some in MR and contralateral DR. Dense labeling in caudal raphe was observed only with hindbrain or spinal cord injections (**Figure S7A**), consistent with previous reports that caudal raphe serotonin neurons preferentially innervate the hindbrain and spinal cord^55,56^.

We next asked whether the spatial distribution of labeled cells in a single animal could predict its projectomic-group identity. We developed a supervised version of NMF^57^ with Bayesian-optimized hyperparameters (*k* = 5; **Methods**), which achieved 83.6% sample-level classification accuracy (5-fold cross-validation; chance = 20%) (**Figure 4C**; **Figure S7B**, **C**). Thus, despite partial overlaps, individual-animal cell distributions reliably reflect projectomic-group identity.

### Transcriptomic and spatial identity jointly define projectomic groups

In well-studied neural systems, such as the vertebrate retina and the fly antennal lobe, single-cell transcriptomic types correspond closely to cell types defined by morphology, connection patterns, and physiological response properties^58–62^. However, the extent to which axonal projection patterns correspond to neuronal transcriptomes in the mammalian brain is still largely an open question^63–67^. We therefore combined retrograde barcode tracing with sequential STARmap^68^ to simultaneously capture the spatial transcriptomic profile and projection target of individual serotonin neurons in the dorsal and median raphe (**Figure 5A**; **Methods**). We designed probes targeting two serotonergic markers (*Tph2* and *Sert*), 10 markers for the 10 transcriptomic clusters of DR and MR serotonin neurons^25^, and five retrograde barcodes (**Figure 5B**; probe sequences in **Table S3**). Fourteen mice received retrograde barcode injections at marker regions of the five projectomic groups (**Table S4**; **Methods**): each of the four “cross-group” mice with injections spanning all five projectomic groups, and ten “within-group” mice with injections confined to a single group (two mice per group).

**Figure 5.**
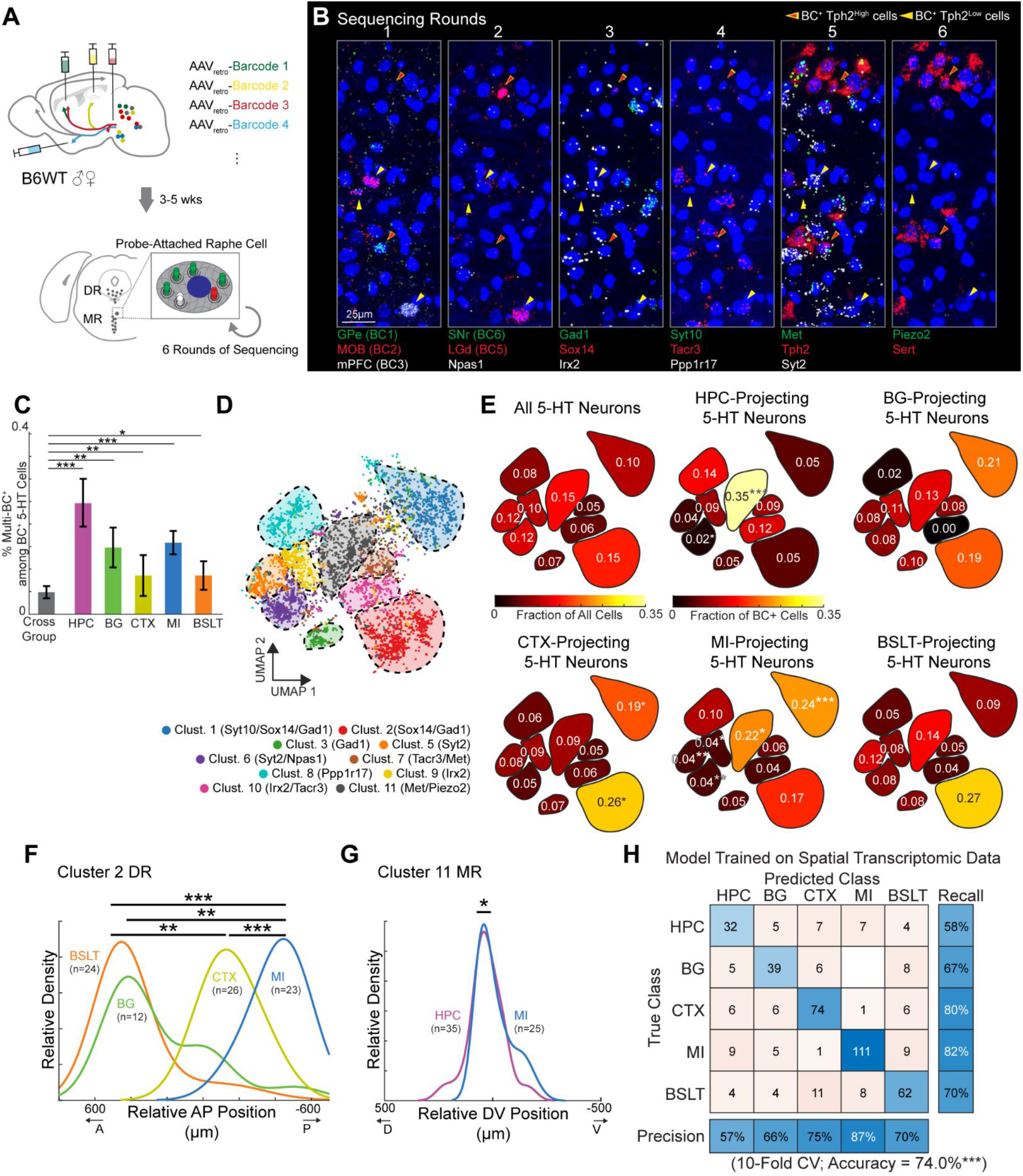
Relationship between transcriptomic clusters, projectomic groups, and cell body locations. **(A)** Six-round sequential STARmap design in the raphe. Probes targeted five barcode sequences, two pan-serotonin markers, and ten transcriptomic cluster markers based on a previous study^25^. **(B)** Example images of the same raphe neurons across sequencing rounds. The first five channels in the first two imaging rounds were assigned to different barcode signals (**Table S3**). Yellow arrows, barcode-positive neurons; red arrows, barcode-positive serotonin neurons. Section thickness: 10 µm; maximal intensity projection. **(C)** Cross- and within-group co-projection rates. Bars, percentage of barcode-positive (BC^+^) cells carrying two or more barcode types; error bars, binomial standard error. n = 227, 61, 65, 35, 70, and 177 cells for the cross-group, HPC-, BG-, CTX-, MI-, and BSLT-groups, respectively. *p < 0.05, **p < 0.01, ***p < 0.001, difference permutation test (50,000 iterations), BKY FDR-corrected. **(D)** UMAP embedding of 6,214 quality-controlled serotonin neurons from the STARmap dataset, colored by 10 transcriptomic subtype as previously described^25^. Shaded regions outline high-density areas for each cluster. **(E)** Transcriptomic composition of barcode-positive neurons for each projectomic group, displayed as cluster masks on the UMAP. Top left, baseline proportions across all serotonin neurons. Remaining panels, fraction of each group’s barcode-positive cells per cluster. Asterisks, significant enrichment or depletion relative to baseline (Fisher’s exact test, BKY FDR-corrected; *p < 0.05, **p < 0.01, ***p < 0.001). **(F)** Anteroposterior distributions of projectomic groups within Cluster 2, DR. Curves smoothed by kernel density estimation. Groups with n < 10 excluded from statistical testing. **p < 0.01, ***p < 0.001, permutation t-test with BKY FDR correction. **(G)** Dorsoventral distributions of HPC- and MI-projecting neurons within Cluster 11, MR. *p < 0.05. **(H)** Confusion matrix from 10-fold cross-validated random forest classifier using combined spatial and transcriptomic features. Class weighting applied for unequal group sizes. n=430 cells; multi-group barcode cells excluded. ***p < 0.001, permutation test, 1,000 iterations (chance 22.0%). See **Figure S8** and **Table S3–S5** for additional data.

Of the 8,715 *Tph2*-positive serotonin neurons identified, 635 (7.3%) were barcode-positive for at least one target (**Table S5**). In within-group mice, 15.2% of barcode-positive neurons (62/408) carried ≥ 2 barcode types—significantly higher than the 4.8% (11/227) in cross-group mice, and the increase was significant in all five groups (**Figure 5C**). Thus, individual serotonin neurons preferentially collateralize within—rather than across—projectomic groups, providing single-cell support for the population-level grouping.

We next mapped serotonin neurons to 10 transcriptomic subtypes using a two-stage consensus classification strategy (75.9% cross-validation accuracy on the reference dataset^25^; **Figure S8A**; **Methods**). Of the 8,715 Tph2-positive neurons, 6,214 (71.3%) reached consensus across four independent classifiers and were retained for subtype analysis—including 439 of the 635 barcode-positive cells (69.1%)—forming distinct UMAP clusters consistent with expected marker expression (**Figure 5D**; **Figure S8B**). To assess associations between transcriptomic identity and projection patterns among these barcode-positive neurons, we computed an enrichment ratio (ER) for each cluster–group pair (**Methods**). Each projectomic group exhibited a distinct molecular signature, defined by a characteristic combination of enriched and depleted subtypes rather than a single subtype (**Figure 5E**; **Figure S8C**): Cluster 1 was enriched in CTX- and MI-projecting serotonin neurons (ER = 1.9 and 2.3, respectively); Cluster 11 in HPC (2.3) and MI (1.5); Cluster 2 in CTX (1.6). The MI group further showed simultaneous depletion of Clusters 5, 6, and 9, and Cluster 6 was also depleted in HPC. As an independent validation, we injected an AAV expressing mCherry in a Cre- and Flp-dependent manner into the raphe of *Piezo2-Cre;Sert-Flp* mice. Most labeled neurons co-expressed *Tph2* (88.2%), and their axons preferentially innervated hippocampal and medial-interbrain regions, consistent with Cluster 11 enrichment in the HPC and MI groups (**Figure S8D–F**). This combinatorial, rather than one-to-one, correspondence between transcriptomic identity and projection target parallels observations in the dopamine^69^ and norepinephrine^70^ systems, where molecularly defined subtypes show overlapping rather than strictly segregated projections.

Beyond transcriptomic composition, projectomic groups were also spatially segregated within shared subtypes. In Cluster 2 (DR), BSLT-projecting serotonin neurons were concentrated anteriorly and MI-projecting neurons posteriorly, with BG- and CTX-projecting neurons in between (**Figure 5F**). In Cluster 11 (MR), HPC-projecting neurons were positioned dorsally and MI-projecting neurons ventrally (**Figure 5G**). These spatial separations suggest that even neurons sharing the same transcriptomic subtype are further organized by projection target within the raphe nuclei.

To quantify the combined contribution at the single-cell level, we trained a Random Forest classifier to predict the projectomic groups of individual barcoded cells from 16 features (10 marker genes, *Tph2*, *Sert*, AP/ML/DV coordinates, DR/MR identity). The combined model achieved 74.0% accuracy (**Figure 5H**), substantially exceeding classifiers trained on transcriptomic (41.4%) or spatial features (58.4%) alone (**Figure S8G, H**). This additive gain suggests that transcriptomic identity and spatial position provide complementary, non-redundant information about projection target.

Several factors likely contribute to the complexity of observed transcriptome–projection correspondence. First, our transcriptomic measurement captures a neuron’s adult gene expression state, which may not fully reflect the developmental programs that guide axon targeting. Prior studies in both *Drosophila*^60,71,72^ and mice^73^ show that transcriptomic differences between closely related neurons are largest during the circuit assembly stage but diminish in adults. If this applies to serotonin neurons, the adult correspondence we observed may underestimate the developmental role of transcriptomic identity. Second, the molecular features most relevant to axonal targeting—guidance receptors and cell adhesion molecules—may not be the genes that distinguish transcriptomic clusters. Cadherins, for example, span multiple clusters yet correlate with cortical projection patterns^74^. The associations we detected may therefore underrepresent the true transcriptome–projection correspondence. Finally, it may be biologically advantageous for the same transcriptomic cluster to innervate multiple target systems, and for multiple transcriptomic clusters to innervate the same target. Different transcriptomic clusters of serotonin neurons can co-release glutamate, GABA, or neuropeptides, and each projectomic group may require multiple clusters to serve distinct functions. For example, Vglut3-expressing serotonin neurons selectively target specific cell types in the basal amygdala, differentially influencing social and anxiety-related behaviors^75^. This logic—where projection target and neurotransmitter identity jointly define a neuron’s function—may explain why the transcriptomic clusters are distributed across projectomic groups and why multiple projectomic groups converge onto the same projection targets.

### Projection-specific Tph2 depletion produces distinct effects across behavioral paradigms

We next investigated the behavioral roles of the five projectomic groups using a loss-of-function approach (**Figure 6A**). We injected AAV_retro_ expressing a Flp-dependent Cre recombinase into a marker region for each group of *Tph2^fl/fl^;Sert-Flp;Ai65* mice (experimental group), depleting Tph2—an enzyme essential for serotonin biosynthesis—specifically in projection-defined serotonin neurons. *Sert-Flp;Ai65* mice with identical virus injections served as controls (no Tph2 depletion). The Flp-gated Cre expression strategy allowed histological labeling of transduced neurons while minimizing Cre toxicity at injection sites. RFP⁺ neurons in the raphe of experimental mice showed near-complete loss of Tph2 immunoreactivity, whereas those in controls retained robust Tph2 expression (**Figure 6B, C**). In target regions, RFP⁺/Sert⁺ axons showed markedly reduced serotonin signal relative to neighboring RFP⁻/Sert⁺ fibers (**Figure 6D**). Across animals, the fraction of Sert⁺ voxels that were RFP⁺ (an indicator of serotonergic axons depleted of Tph2) was substantially greater in target than in non-target regions (**Figure 6E**). Together, these data showed effective projection-specific depletion of Tph2 and serotonin production.

**Figure 6.**
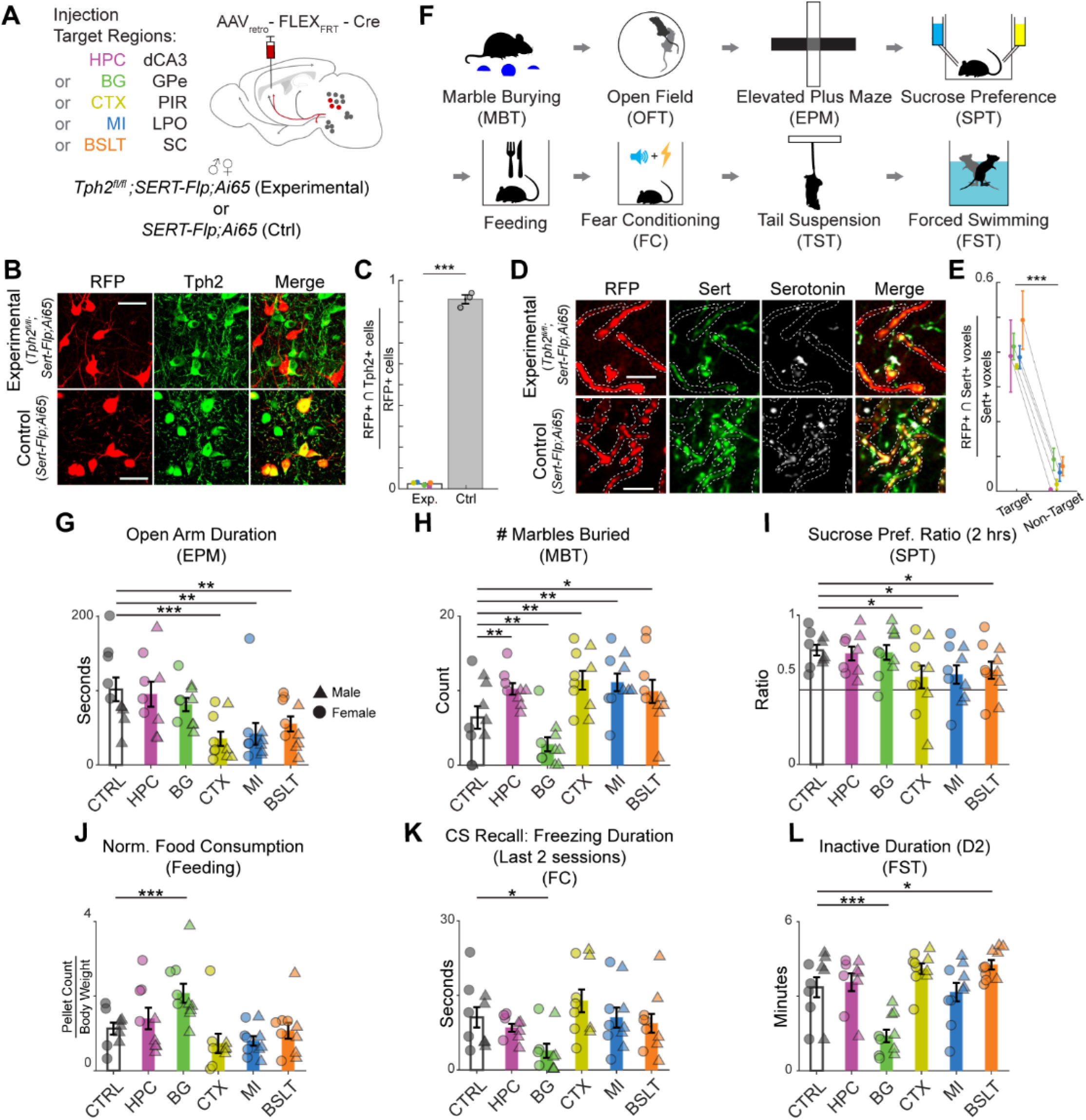
Projection-specific *Tph2* cKO mice show distinct behavioral profiles compared to controls. **(A)** Experimental paradigm. Virus injections were performed in both males and females for each experimental group (n = 5 males, 4 females in dCA3; 6, 4 in GPe; 4, 6 in PIR; 5, 5 in LPO; 5, 5 in SC). For the control group, one male and one female were injected for each of the five target regions (total n = 5 males, 5 females). Abbreviations: dCA3, dorsal CA3; GPe, globus pallidus externus; PIR, piriform cortex; LPO, lateral preoptic area; SC, superior colliculus. **(B)** Tph2 immunoreactivity in RFP⁺ raphe neurons from experimental (*Tph2^fl/fl^;Sert-Flp;Ai65*) and control (*Sert-Flp;Ai65*) animals. Scale bar 50 µm. **(C)** Fraction of RFP⁺ neurons co-expressing *Tph2*. Each dot, one animal (Exp, n = 5; Ctrl, n = 3, one each injected in dCA3, GPe, and PIR); colors in Exp follow panel A. Bars, mean ± SEM. ***p < 0.001, bootstrap test on difference of means (10,000 iterations). **(D)** Sert, RFP, and serotonin immunostaining in target regions; RFP⁺/Sert⁺ axons show reduced serotonin relative to RFP⁻/Sert⁺ fibers. White dashed lines, RFP⁺ axons. Scale bar 10 µm. **(E)** Fraction of Sert⁺ voxels co-labeled with RFP in target (two marker regions of the injected group) vs. non-target (eight marker regions of the other four groups), averaged per animal (mean ± SEM). Each color, one animal (color scheme as in panel A); lines, paired measurements (n = 5). ***p < 0.001, paired bootstrap test of the mean difference (10,000 iterations) **(F)** Sequence of behavioral tests. **(G–L)** Selected behavioral effects caused by disrupting serotonin synthesis in specific projectomic groups. Two-way ANOVA followed by pairwise bootstrap tests comparing each experimental group to control (5 comparisons, 200,000 iterations) with BKY FDR correction. Sexes pooled within groups when the sex × group interaction was not significant (*p*_interaction_ > 0.05). *p < 0.05; **p < 0.01; ***p < 0.001. See **Figure S9, S10** and **Table S6** for additional data.

Next, we assessed behavior across eight paradigms (**Figure 6F**) commonly used to assay the function of serotonergic neurons and their potential relevance to human psychopathology. Tests were ordered by ascending stress levels to minimize confounds due to cumulative stress. These included the marble burying test^76–78^ (MBT), four-phase open field test (OFT)^79^, elevated plus maze (EPM)^80–82^, sucrose preference test (SPT)^83,84^, feeding test^85,86^, fear conditioning (FC), tail suspension test^83,87^ (TST), and two-day forced swim test (FST)^24,84^. Video recordings were quantified using MoSeq^88^ (for OFT) and a keypoint-free, SlowFast^89^-based behavior classifier (for MBT, EPM, FC, TST, FST), yielding 54 features across all eight behavioral paradigms (**Figure S9**; **Table S6**; **Methods**). Below, HPC, BG, CTX, MI, and BSLT groups refer to Tph2 depletion in serotonin neurons retrogradely labeled via AAV_retro_ injection into their respective marker regions, dCA3, GPe, PIR, LPO, and SC (**Figure 6A**; **Figure S4C**).

Using these behavioral assays, we identified effects of projection-specific serotonin deprivation across multiple domains. In the elevated plus maze, the CTX, MI, and BSLT groups showed reduced open arm exploration compared to controls (**Figure 6G; Figure S10A**), a measure that has been interpreted as reflecting anxiety-related behavior^80–82^. In the marble burying test, the HPC, CTX, MI, and BSLT groups buried significantly more marbles (**Figure 6H**). The co-occurrence of increased marble burying and reduced open-arm exploration in CTX, MI, and BSLT groups suggests that serotonin deprivation in these projection targets promotes both anxiety-related and repetitive behaviors.

The CTX, MI, and BSLT groups also showed a significant reduction in sucrose preference over water (two-hour session; **Figure 6I**), with no significant differences in total volume consumed (**Figure S10C**), indicating that the reduced sucrose preferences were not attributable to overall fluid intake. As reduced sucrose preference is commonly interpreted as reflecting a shift in reward-related behavior^83,84^, its co-occurrence with increased anxiety-related and repetitive behaviors within these groups suggests that multiple projectomic groups jointly affect anxiety-related, repetitive, and reward-related behaviors.

The BG group was particularly distinctive, diverging from other groups across several behavioral domains. The strong reduction of marble burying (**Figure 6H**) but no effect in elevated plus maze (**Figure 6G**) may suggest that serotonergic projection to BG preferentially promotes components of repetitive behavior, consistent with previously reported involvement of the basal ganglia in obsessive/compulsive-like behavior^90,91^. The BG group was also the only group to show significantly increased food consumption after 24-h food deprivation (**Figure 6J**). Likewise, it uniquely showed reduced freezing in the final two recall sessions of fear conditioning, suggestive of faster fear extinction (**Figure 6K**; **Figure S10D–F**). It was also the only group with reduced immobility in forced swim test, commonly interpreted as reflecting increased active coping (**Figure 6L**)^24,87^. These data suggest that serotonergic modulation of basal ganglia inhibits overeating, slows fear memory extinction, and suppresses active coping in the face of adversity.

Notably, we observed pronounced sex differences in certain combinations of manipulation and behavioral paradigm. For example, control and BSLT females spent more time in the open arm than male counterparts (**Figure 6G**; **Figure S9B**, "Open Arm Duration"), whereas MI and BSLT males showed greater immobility in forced swim test on both days ("Inactive Duration, Day 1 and Day 2"). Thus, serotonergic projections can regulate behavior in a sex-dependent manner, with the same projection targets producing distinct—and sometimes opposite—outcomes in males versus females (**Figure S9B**).

Taken together, our systematic loss-of-function approach reveals that each projectomic group affects behaviors across multiple paradigms, rather than within paradigms tied to a single behavioral interpretation. At the same time, each group produced a distinct combination of effects with different magnitudes, indicating that projectomic identity shapes a specific behavioral profile. The complete dataset—effect sizes (Cohen’s *d*), sex comparisons, and statistics for all 54 features—is provided in **Figure S9** and **Table S6** as a resource for independent reanalysis and cross-study comparison.

### Serotonergic projectomic groups organize the behavioral landscape into distinct domains

Behavioral phenotypes are multifaceted: shifts in one behavior are often accompanied by changes in others (**Figure 6**). Examining this covariance structure may provide a more complete view than single features in isolation, capturing both broad patterns and subtle differences. We therefore asked whether the five projectomic groups occupy distinct positions within a shared behavioral latent space. To define this space, we applied multi-class linear discriminant analysis^92^ (LDA) to find the combinations of behavioral features that best separate predefined groups (**Figure 7A**; **Methods**). We trained the model on samples from the dCA3-, GPe-, PIR-, LPO-, and SC-injected groups (**Figure 6**), analyzing males and females separately given the pronounced sex differences observed in a small subset of univariate analyses (**Figure 6G**; **Figure S9**). Cross-validation yielded classification accuracies of 79% for females and 60% for males (chance level from permutation test: females 25.9 ± 7.2%, males 26.1 ± 7.0%; **Figure S11**; **Methods**), indicating that serotonin depletion in distinct projectomic groups produces characteristic behavioral signatures. Projecting all samples onto the full-model LDA space further confirmed clear group separation, with same-site animals clustering together (**Figure 7B, C**).

**Figure 7.**
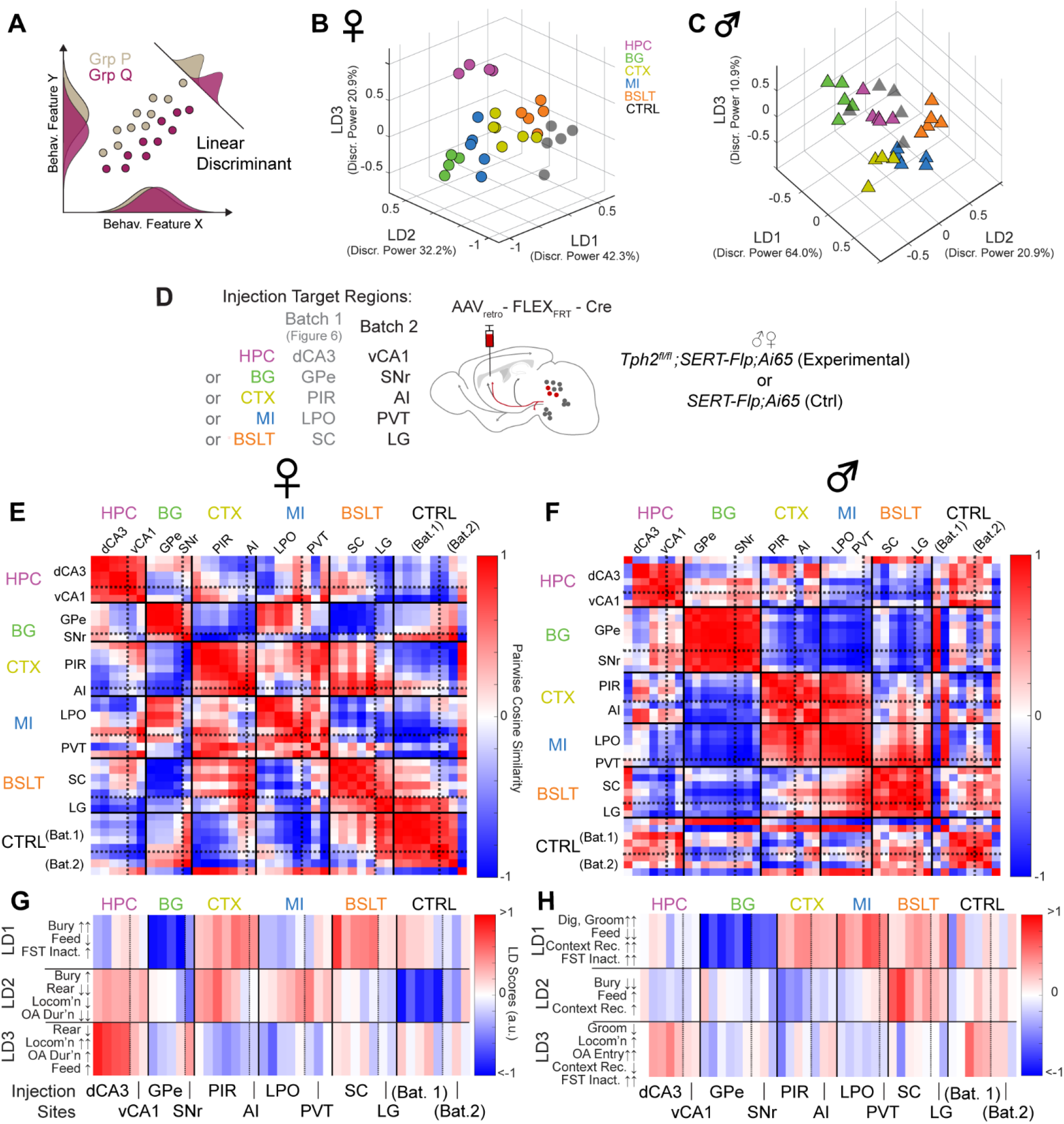
Multidimensional behavioral phenotyping of projection-specific *Tph2* conditional knockout mice using LDA and cross-batch validation. **(A)** Rationale for using linear discriminant analysis (LDA). Although hypothetical groups P and Q exhibit substantial overlap in individual feature dimensions X and Y, projecting them onto a linear discriminant axis results in clear separation in a lower-dimensional space. **(B, C)** Samples in Figure 6 were plotted in the first three LD axes for females (**B**) and males (**C**). LDs were derived to maximize separation among the six groups. Discriminant power, each axis’s share of total group separation (sums to 1 across all LDs; analogous to "variance explained" in PCA); Dots, individual animals; colors, group identity. **(D)** Batch 2 injection sites: vCA1 (HPC; n = 2 females, 2 males), SNr (BG; 1, 3), AI (CTX; 1, 3), PVT (MI; 3, 1), and LG (BSLT; 2, 2), along with 3 control mice per sex. The LDA model was trained exclusively on Batch 1, and Batch 2 samples were then projected onto the same latent space. Abbreviations: vCA1, ventral CA1; SNr, substantia nigra reticulata; AI, agranular insular cortex; PVT, paraventricular nucleus of the thalamus; LG, lateral geniculate nucleus. **(E, F)** Pairwise cosine similarity between samples, computed using their LD scores in the Batch 1–trained latent space. **(G, H)** LD scores of Batch 1 and Batch 2 samples. Axis interpretations (left of colormaps) based on LD loadings and structure coefficients described in **Figure S11D–G**. Same sample order as panels E and F, respectively. See **Figure S11** and **S12** for additional data.

Given the projectomic organization of serotonin neurons, we next tested whether serotonin depletion in each projectomic group produces a characteristic behavioral phenotype, independent of the specific marker region targeted. We generated an independent dataset using a different marker region per group (hereafter Batch 2, with the original training set referred to as Batch 1; **Figure 7D**). Individual Batch 2 samples were projected onto the latent space trained exclusively on Batch 1 (with covariance alignment^93^ between batches; **Methods**), enabling us to test whether the discriminative structure defined by the original training set generalized to independent marker regions. To test whether samples sharing the same projectomic group resemble each other regardless of marker region or batch, we computed pairwise cosine similarities across all Batch 1 and Batch 2 samples. We found that samples injected within the same projectomic group formed distinct high-similarity patches along the diagonal, both within each batch and across Batch 1 and Batch 2 despite their different marker regions, confirming within-group consistency (**Figures 7E**, **F**). Furthermore, within-group Batch 1–Batch 2 similarities significantly exceeded between-group similarities in all groups in both sexes (**Figure S12**). The only exception was the control group; it is possible that in the absence of a shared manipulation, individual variability may dominate any group-level pattern, explaining why within-group similarity for the control group did not exceed between-group similarity (**Figure S12**). These results demonstrate that the observed behavioral phenotypes are organized by projectomic identities rather than specific injection sites.

Having established the cross-batch reproducibility of within-group structure, we next examined the composition of each behavioral latent axis (**Figures 7G**, **H**; full model details in **Figure S11D–G**). LD1 in both sexes—despite being derived independently—included an overlapping set of behavioral features, such as increased marble burying/digging, decreased feeding, and increased immobility in forced swim test, suggesting that these measures capture salient axes of serotonergic behavioral variation independent of sex. LD2 and LD3, by contrast, diverged between sexes. In females, LD2 reflected increased marble burying alongside decreased locomotion, open-arm duration, and rearing, whereas LD3 captured decreased rearing alongside increased locomotion, open-arm duration, and feeding. In males, LD2 reflected decreased marble burying alongside increased feeding and late-stage contextual fear recall, whereas LD3 captured decreased grooming and late-stage contextual fear recall alongside increased locomotion, open-arm entries, and immobility in forced swim test. These differences suggest sex-specific dimensions of serotonergic regulation, with the primary discriminative axis (LD1) anchored by largely sex-shared behavioral features while subsequent axes (LD2, LD3) diverge between males and females.

At the individual sample level, Batch 2 samples projected into the same Batch 1–derived latent space showed LD scores consistent with their projectomic groups (**Figure 7G, H**). BG-injected groups showed the most negative LD1 scores in both sexes, associated with reduced marble burying and immobility in forced swim test alongside increased food intake (**Figure 6H, L,** and **J**). In females, CTX and BSLT groups overlapped on LD1 and were primarily separated by LD2, with the CTX group characterized by reduced feeding, open-arm duration, and locomotion (**Figure 6J**, **G**; **Figure S10G**). HPC-injected females exhibited the highest LD3 scores among all groups, reflecting increased feeding, locomotion, and decreased rearing (**Figure 6J**; **Figure S10G**, **H**). In males, CTX, MI, and BSLT groups similarly overlapped on LD1, reflecting their behavioral similarity across feeding, immobility in forced swim test, and grooming (**Figure 6J**, **L**; **Figure S10B**), with BSLT males further separated along LD2 by reduced marble burying (**Figure 6H**). Notably, despite this shared LD1 structure across sexes, the MI group projected onto LD1 in opposite directions in males and females, suggesting sex differences for serotonergic modulation of MI-projection targets. Several of these distinctions were not apparent in individual measures but emerged only when behaviors were considered in their covariant structure, demonstrating how multivariate analysis can capture phenotypes obscured by univariate analyses alone.

### Projection-defined subsystems shape behavioral phenotypes

To our knowledge, the projectomic organization described here is consistent with previously published studies perturbing serotonin neurons projecting to specific brain regions. Depleting serotonin from serotonergic projections to the central amygdala, a marker region within the BG group, produced reduced negative-valence behaviors^24^, consistent with our GPe- and SNr-injected animals. Suppressing projections to the lateral habenula (MI-group marker region) reduced sucrose preference without affecting immobility in forced swim test in male rats^94^, consistent with our LPO- and PVT-injected male animals. Disrupting serotonergic input to the cerebellar cortex (BSLT marker region) elicited anxiety-related behavior in male mice^95^, paralleling observations in our SC- and LG-injected animals. These convergent findings, despite varied experimental conditions, reinforce the notion that the five projectomic groups represent functionally coherent units of serotonergic behavioral modulation.

Beyond group-level dissociations, our analyses also revealed notable instances of phenotypic overlap among projectomic groups targeting distinct brain regions. This may reflect the partial target sharing between groups or convergence onto common downstream circuit nodes. The marker regions and projectomic framework defined here provide entry points to distinguish these possibilities.

The co-occurrence of negative-valence phenotypes we observed is reminiscent of the frequent comorbidity of depressive, obsessive-compulsive, and anxiety disorders in humans, despite their categorical distinction in clinical settings^96,97^. In our data, depleting serotonin in a single projectomic group was sufficient to alter behaviors across multiple paradigms, such as elevated plus maze, marble burying, and sucrose preference. Disruption of different projectomic groups, in turn, produced distinct combinations of these effects. Together, these observations suggest a circuit basis for understanding such psychiatric conditions as overlapping continua of symptoms rather than discrete categories^98,99^. By decomposing co-occurring behavioral phenotypes into quantifiable dimensions, our approach may help generate circuit-level hypotheses for understanding clinical heterogeneity.

### Limitations of the study

First, our projectome mapping is based on bulk axon tracing of a group of serotonin neurons projecting to the same target sites. This approach may overlook heterogeneity of projections within the projectomic groups, as suggested by subpatterns we identified. Future tracing of axonal projections of individual serotonin neurons^25^ but at a much larger scale, analyzed in a manner similar to this work, could elucidate the serotonin projectome at a finer resolution. Second, since our analyses are contingent on the dataset used, we cannot rule out the possibility that additional projectomic groups and particularly subgroups within each group could emerge when data from samples injected into more regions are included. Third, the intersectional reporter mouse line^29^ for tracing whole-brain axonal projection independent of cell body position utilizes a cytosolic marker that does not distinguish between synaptic terminals and axons-in-passage. Although we attempted to reduce the influence of axons-in-passage by excluding their density in the fiber tracts defined in CCFv3, we cannot exclude axons near the terminals from contributing to our projectome analysis. Future studies employing a presynaptic terminal marker could enable more precise identification of serotonergic innervation patterns. Finally, our behavioral characterization used projection-specific *Tph2* deletion as a loss-of-function approach; we did not examine gain-of-function effects or perform in vivo physiological recordings of the targeted projectomic populations. These complementary studies—utilizing similar marker regions and analytical pipelines described in this study—will help further refine how serotonergic signaling in each projectomic group shapes the behavioral phenotypes we report here.

## CONCLUDING REMARKS

Our work demonstrates that projectomic identity—the whole-brain pattern of axon collateralization—provides a powerful framework for decomposing the serotonin system into its functional, constituent parts. Combining whole-brain axon tracing, spatial transcriptomics with retrograde barcoding, and behavioral analysis using projection-specific serotonin loss-of-function, we identified five projectomic groups for DR and MR serotonin neurons and revealed three organizational principles. First, serotonin neurons preferentially co-innervate functionally related brain regions rather than physically adjacent ones. Second, projectomic identity emerges from a combination of transcriptomic subtype and spatial position of serotonin neurons. Third, projection-specific serotonin depletion produces dissociable behavioral phenotypes that generalize across independent marker regions within each projectomic group. Our projectomic, spatial transcriptomic, and behavioral data— as well as the analysis pipelines—provide a resource for future anatomical, physiological, and behavioral studies of the serotonin system.

## RESOURCE AVAILABILITY

### Lead Contact

Further information and requests for resources and reagents should be directed to the lead contact, Liqun Luo.

### Materials availability

All unique reagents generated in this study are available from the lead contact.

### Data and code availability

- The raw image files are available on Brain Image Library (https://doi.org/10.35077/g.1213)
- Data analysis code and relevant data can be accessed at https://github.com/junhsong92/serotonin_projectome.
- Any additional information is available upon requests from the corresponding author.

## ACKNOWLEDGEMENT

We thank members of the Luo lab for their constructive feedback on this project and manuscript, especially Alex Starr, David Wang, Lijun Qi, and Chloe Bair-Marshall. We thank Larry Swanson, Joel Hahn, Justus Kebschull, Neir Eshel, Robert Malenka, Daniel Cardozo-Pinto, and Wendy Wenderski for critiques on the manuscript. We thank Xiao Wang and Hailing Shi for sharing circular barcode constructs for STARmap and Mary Molacavage for administrative assistance. STARmap imaging was performed at the Stanford University Cell Sciences Imaging Facility (CSIF) (RRID:SCR_017787). J.H.S. is supported by the Bio-X Fellowship and the Paul and Mildred Berg Graduate Fellowship in Biology. L.L. is an investigator of the Howard Hughes Medical Institute. This work was supported by BRAIN Initiative grants from the National Institutes of Health (R01-NS104698 to L.L.) and (R01-NS131987 to L.L. and S.W.L.).

## AUTHOR CONTRIBUTIONS

J.H.S., D.F., and L.L. conceived this project. J.H.S. designed and conducted the main data analyses. J.H.S., D.F., A.M., I.R., Q.W., and S.A.X. prepared samples for axon tracing. J.H.S. and D.F. preprocessed whole-brain imaging data. J.H.S. and Y.W. performed STARmap experiments and analyses. J.H.S., I.R., J.K.B., and K.Z. performed behavioral analyses. J.H.S., K.Z., and T.H.S. preprocessed behavioral recordings. J.H.S. and Y.Y. produced viruses. X.C. provided AAV_8retro_ capsid plasmid. S.W.L. advised on data analyses. J.H.S. and L.L. wrote the manuscript with input from all co-authors. L.L. supervised the work.

## DECLARATION OF GENERATIVE AI AND AI-ASSISTED TECHNOLOGIES IN THE WRITING PROCESS

The first author used Claude Opus 4.8 (Anthropic) to correct grammatical errors and typos. The authors thoroughly reviewed and edited Claude’s outputs and take full responsibility for the published article.

## MATERIALS AND METHODS

### KEY RESOURCE TABLE

| REAGENT or RESOURCE | SOURCE | IDENTIFIER |
| --- | --- | --- |
| **Antibodies** |  |  |
| anti-RFP rabbit polyclonal antibody | Rockland | Cat# 600-401-379 |
| anti-tdTomato goat polyclonal antibody | Origene | AB8181-200 |
| anti-Tph2 goat polyclonal antibody | Abcam | ab121013 |
| anti-SERT guinea pig polyclonal antibody | Nittobo Medical | HTT-GP-Af1400 |
| anti-Serotonin rabbit antibody | Immunostar | 20080 |
| **Bacterial and Virus Strains** |  |  |
| AAV<sub>8retro</sub>-CAG-FLEX<sup>loxP</sup>-Flp | Custom Packaged; Capsid construct from Wang et al., 2026<sup>100</sup>; Ren et al., 2018<sup>24</sup> | N/A |
| AAV<sub>8retro</sub>-U6-circBC(1-6)-hSyn-V5-2A-Flag-WPRE-pA | UNC Neurotools; capsid plasmid from Wang et al., 2026<sup>100</sup>; Shi et al., 2023<sup>101</sup> | N/A |
| AAV<sub>8retro</sub>-CAG-FLEX<sup>FRT</sup>-Cre | Custom Packaged; Capsid plasmid from Wang et al., 2026<sup>100</sup>; Construct modified from Addgene 121675 | N/A |
| AAV-Con/Fon-mCherry | Addgene | 137132-AAV8 |
| **Chemicals and Reagents** |  |  |
| Red retrobeads | LumaFluor Inc. |  |
| Gel Slick Solution | Lonza | 50640 |
| Bind-Silane | SERVA | 28739.01 |
| Poly-L-lysine solution, 0.1% w/v | Sigma | P8920 |
| Ultrapure distilled water | Invitrogen | 10977-015 |
| Glass bottom 24-well | MatTek | P24G-1.5-13-F |
| #2 Micro coverglass, 12 mm diameter | Electron Microscope Sciences | 72226-01 |
| O.C.T. Compound | Fisher scientific | 23-730-571 |
| 16% PFA, EM grade | Electron Microscope Sciences | 15710-S |
| Methanol for HPLC | Fisher scientific | A412SK-4 |
| PBS, 7.4 | Thermo Fisher Scientific | AM9625 |
| Tween-20, 10% solution | Calbiochem | 655206 |
| Triton-X-100, 10% solution | Millipore Sigma | 93443 |
| Deionized Formamide | Millipore Sigma | 344206-1L |
| 20x SSC buffer | Millipore Sigma | S6639 |
| Salmon sperm DNA | Invitrogen | AM9680 |
| SUPERasein | Invitrogen | AM2696 |
| T4 DNA ligase | Thermo Fisher Scientific | EL0011 |
| Phi29 DNA polymerase | Thermo Fisher Scientific | EP0094 |
| 10 mM dNTP mix | Invitrogen | 100004893 |
| BSA, molecular biology grade | New England Biolabs | B9000S |
| 5-(3-aminoallyl)-dUTP | Invitrogen | AM8439 |
| Acrylic acid NHS ester, 90% | Millipore Sigma | A8060 |
| DMSO, anhydrous | Molecular Probes | D12345 |
| Acrylamide solution, 40% | Bio-Rad | 161-0140 |
| Bis Solution, 2% | Bio-Rad | 161-0142 |
| Ammonium persulfate | Millipore Sigma | A3678 |
| N,N,N,N-Tetramethylethylenediamine | Millipore Sigma | T9281 |
| OminiPur SDS, 20% | Calbiochem | 7991 |
| Protease K, RNA grade | Invitrogen | 25530049 |
| Benzyl Ether | Millipore Sigma | 108014 |
| Ethyl Cinnamate | Millipore Sigma | 112372 |
| 20 mg dustless precision pellets | Bio-Serv | FF-0071 |
| Peppermint oil | Sigma-Aldrich | W284807 |
| Artificial garlic oil | Sigma-Aldrich | W530316 |
| **Oligonucleotides** |  |  |
| STARmap probes for the marker genes and barcodes (See **Table S3**) | Integrated DNA Technology | N/A |
| **Experimental Models: Organisms/Strains** |  |  |
| Mouse: Tg(Slc6a4-cre)ET33Gsat/Mmucd (Sert-Cre) | MMRRC | MMRRC, Stock #017260-UCD |
| Mouse: Tg(Tph2-icre/ERT2)6Gloss/J (Tph2-iCreER) | The Jackson Laboratory | JAX: 016584 |
| Mouse: B6;129S-Gt(ROSA)26Sortm65.1(CAG-tdTomato)Hze/J (Ai65) | The Jackson Laboratory | JAX: 021875 |
| Mouse: B6.Cg-Slc6a4tm1.1(flpo)Luo/J (Sert-Flp) | The Jackson Laboratory | JAX: 034050 |
| Mouse: B6(SJL)-Piezo2tm1.1(cre)Apat/J (Piezo2-Cre) | The Jackson Laboratory | JAX: 027719 |
| Mouse: B6(Cg)-Crhtm1(cre)Zjh/J (Crh-Cre) | The Jackson Laboratory | JAX: 012704 |
| Mouse: C57BL/6J | The Jackson Laboratory | JAX: 000664 |
| Mouse: Tph2tm1Msc1 (Tph2flox/flox) | Wu et al., 2012<sup>102</sup> | N/A |
| **Software and Algorithms** |  |  |
| IMARIS | Bitplane | https://www.bitplane.com/ |
| MATLAB | Mathworks | https://www.mathworks.com/ |
| Custom MATLAB script for NMF-based axon analysis | This paper | https://github.com/junhsong92/serotonin_projectome |
| Custom MATLAB script for supervised-NMF analysis | This paper | https://github.com/junhsong92/serotonin_projectome |
| Custom Python script for behavior classification | This paper | https://github.com/junhsong92/serotonin_projectome |
| Ilastik | Berg et al., 2019<sup>103</sup> | https://www.ilastik.org/ |
| Elastix | Klein et al., 2010<sup>104</sup> | https://elastix.dev/ |
| Trailmap | Friedmann et al., 2020<sup>33</sup> | https://github.com/albert597/TRAILMAP |
| Cellpose | Stringer et al., 2021<sup>105</sup> | https://www.cellpose.org/ |
| Baysor | Petukhov et al., 2021<sup>106</sup> | https://github.com/kharchenkolab/Baysor |
| Allen CCFv3 | Allen Institute for Brain Science<sup>34</sup> | https://download.alleninstitute.org/informatics-archive/current-release/mouse_ccf/ |
| AMaSiNe | Song et al., 2020<sup>107</sup> | https://github.com/cogilab/AMaSiNe |
| Harmony/Harmonypy | Korsunsky et al., 2019<sup>108</sup> | https://github.com/slowkow/harmonypy |
| Scanpy | Wolf et al., 2018<sup>109</sup> | https://scanpy.readthedocs.io/en/stable/ |
| umap-learn / densMAP | Narayan et al., 2021<sup>110</sup> | https://umap-learn.readthedocs.io/en/latest/ |
| MoSeq2 | Wiltshko et al., 2020<sup>88</sup> | https://github.com/dattalab/moseq2-app |
| SlowFast network | Feichtenhofer et al., 2019<sup>89</sup> | https://github.com/facebookresearch/SlowFast |
| YOLOv8 | N/A | https://github.com/ultralytics/ultralytics |
| CORAL | Sun et al., 2016<sup>93</sup> |  |
| **Hardware** |  |  |
| Ultramicroscope II light-sheet microscope | Miltenyi Biotec |  |
| Blaze light-sheet microscope | Miltenyi Biotec |  |
| Nikon Eclipse spinning disk microscope | Nikon |  |
| Microsoft Kinect depth camera | Microsoft |  |
| Bpod State Machine r2 | Sanworks | Product ID 1024 |
| Infrared lick ports | Sanworks | Product ID 1004 |
| FED3.2 pellet dispenser | Open Ephys |  |
| Fear conditioning chamber | Coulbourn Instruments |  |
| HD webcam (C920x HD Pro) | Logitech | C920x |

#### Mouse model

All animal procedures followed animal care guidelines approved by Stanford University’s Administrative Panel on Laboratory Animal Care. All mice were housed on a 12-hour light (7 am to 7 pm)/dark cycle (7 pm to 7 am), and food and water were provided ad libitum. For most of the anatomical experiments conducted (**Figures 1 to 4**), mice aged between 6 and 14 weeks were used. Spinal cord injections were performed on postnatal day 1.5–2.5. The *Ai65* reporter (JAX #021875; C57BL/6J background), *Sert-Cre* (MMRRC, Stock #017260-UCD; C57BL/6J background), and *Tph2-iCreER* (JAX #016584; C57BL/6NTac and C57BL/6J mixed background) were used. *Sert-Cre;Ai65* mice were used for most experiments except for cortex-, medial- and lateral geniculate-injected brains, where *Tph2-iCreER;Ai65* mice were used (as non-specific labeling of cells was observed near some cortex-, medial-, and lateral geniculate body-injection sites in *Sert-Cre;Ai65* mice). The *Tph2-iCreER;Ai65* mice were i.p. injected with 50 mg/kg 4-hydroxytamoxifen (4-OHT; Sigma, Cat# H6278) three weeks after virus injection. For the spatial transcriptomic experiments (**Figure 5**), C57BL/6J wild-type mice aged between 6–8 weeks were used. For the behavioral experiments (**Figures 6, 7**), *Tph2^fl/fl^;Sert-Flp;Ai65* (experimental) and *Sert-Flp;Ai65* (control) mice aged 6–8 weeks at injection were used. Both male and female mice were used for all experiments and data were analyzed together unless stated otherwise.

#### Sample preparation for whole-brain imaging

For virus injections into the brain, *AAV8_retro_-CAG-FLEx-Flp* (titer = 5.0e13 viral genomes (vg)/ml; custom packaged)^24^, mixed with red Retrobeads (1:25 dilution, Lumafluor, Red Retrobeads; for marking injection centers), was injected into the specified regions of the right hemisphere (coordinates and volumes listed in **Table S1**) of male and female mice. Mice were prepared across seven experimental batches. The method for spinal cord injections was adapted from a previous study^111^. Eight weeks after injection, the brains were collected. Mice were first anesthetized with Avertin, then perfused transcardially with phosphate-buffered saline (PBS) (AM9625, ThermoFisher) solution containing heparin at a final concentration of 10 µg/ml (diluted 1:1000), followed by perfusion with ice-cold 4% paraformaldehyde (PFA) (15710-S, Electron Microscope Sciences). After sample collection, all brains were post-fixed overnight at 4°C in 4% PFA in PBS and rinsed three times in PBS the next morning. The brains were subsequently processed using the previously described AdipoClear immunolabeling protocol^32,33^. The entire process for clearing and staining took 28 days, including an 11-day incubation with the primary antibody against RFP (1:500, 600-401-379, Rockland) and a 7-day incubation with secondary antibody (donkey anti-rabbit AlexaFluor 647, ThermoFisher Scientific). Prior to use, all antibody solutions were centrifuged at 12,000 rpm and 20°C for 30 minutes.

#### Light-sheet image acquisition

The general procedure follows the approach outlined in a previous report^33^. In short, more than 24 hours after clearing, brain samples were imaged on a light-sheet microscope (Ultramicroscope II or Blaze, Miltenyi Biotec) in either dibenzyl ether (Millipore Sigma, 108014) or ethyl cinnamate (Millipore Sigma, 112372) to match the refractive index. Samples were imaged using 488 nm, 561 nm, and 647 nm excitation wavelengths to visualize tissue autofluorescence, injection sites marked by Red Retrobeads, and stained axons, respectively. Due to the working depth of the objective used with the Ultramicroscope II, imaging covered the entire right hemisphere (ipsilateral to the injection sites) but only the medial part of the left hemisphere. The full imaging window closely resembles the view depicted in **Figure S1B-K**. For axon imaging, the primary channel (647 nm) was acquired at a 3-μm *z*-step size using the continuous light-sheet scanning method with a contrast-adaptive algorithm (20 acquisitions per plane, numerical aperture 0.1); the 488 nm and 561 nm channels were imaged without dynamic focusing. The final voxel resolutions were 4.1 × 4.1 × 3 μm and 5.9 × 5.9 × 3 μm for images obtained with the Ultramicroscope II and Blaze, respectively. No microscope-dependent differences in quantification results were observed.

#### Light-sheet image pre-processing

Two segmentation models were trained: one specifically optimized for axon segmentation and the other for raphe cell body segmentation. The same signal channels (namely, the 647-nm channel for tdTomato staining) were processed twice using these models with TrailMap^33^. Following visual inspection of the segmentation results, raw image windows with inadequate segmentation were cropped, re-processed with Ilastik^103^, and substituted. Segmented image volumes were then registered to the Allen Reference Atlas CCFv3 (10 μm^3^ voxel resolution upscaled to 5 μm^3^) following previously described methods^33^. In short, the 488-nm autofluorescence image was matched to the reference atlas by applying a series of linear and non-linear transformations using Elastix^104^. Transformation parameters obtained from these processes were then utilized to warp the segmented axon and cell body data to align with CCFv3. The CCFv3-registered segmented axon data were further processed based on the probabilistic volumes of CCFv3-registered segmented axons, as described in a previous study^33^. Briefly, the probabilistic volumes were binned into eight levels (ranging from 0.2 to 0.9), skeletonized, and summed with probability-threshold weighting. To visualize sample brains from an oblique angle, as shown in **Figure 1C**, we used Imaris (Bitplane) to create 3D renderings of the CCFv3 "outer shell" volume with overlaid axon data. Visualizations in other figures were generated using custom-built MATLAB scripts.

#### Axon data analysis using unsupervised NMF

All analyses were performed in MATLAB R2021b (MathWorks). Regional axon data, representing either local density or volume (quantity), were computed by summing the regional voxel values in the processed CCFv3-registered axon volume (for quantity) and dividing by the regional volume (for density). In a small number of injected brains (IDs 71–74; *Sert-Cre;Ai65*), we observed fewer than 10 labeled cell bodies at the injection sites in the ventral posteromedial nucleus of the thalamus (VPM) and the dorsal lateral geniculate nucleus (LGd), likely reflecting transient *Sert* expression. Because these cells were confined to sensory thalamic and primary cortical regions, axon quantification values were set to 0 in those regions to remove confounding signal. The resulting matrix (110 brain samples × 280 regions) was L2-normalized across brain samples and is referred to as ***X***.

As illustrated in **Figure 2B**, a "speckled" cross-validation procedure was used: 90% of the elements of ***X***, randomly scattered across rows and columns, were assigned to ***X_train_***, and the remaining 10% to ***X_test_***. NMF was performed on ***X_train_***across integer values of k from 1 to 15, and reconstruction error on ***X_test_*** was recorded as RMSE_Test_. The procedure was repeated 1,000 times per k value to obtain stable estimates. To benchmark NMF performance, we additionally computed two reference baselines. RMSE_Replicate_ (theoretical performance ceiling) was estimated as the RMSE between a held-out brain sample and the mean of its biological replicates (other brains injected at the same site); this leave-one-out procedure was applied only to injection sites with n ≥ 2 replicates and was repeated 1,000 times. RMSE_Shuffle_ (chance-level floor) was computed by training NMF on a randomly shuffled version of ***X_train_*** and evaluating reconstruction error on the original (unshuffled) ***X_test_***, also repeated 1,000 times per k. RMSE_Test_ values were compared across k values using one-way ANOVA followed by Bonferroni correction. The optimal k was selected as the smallest value at which RMSE_Test_ reached the RMSE_Replicate_ ceiling and was clearly separated from the RMSE_Shuffle_ floor (**Figure 2C**). With k fixed, NMF was repeated 5,000 times with random initializations, and the factorization with the lowest reconstruction error was retained for downstream analyses. The proportion of variance explained by the final factorization was computed as 1 − ||***X*** − ***WH***||^2^*_Fro_* / ||***X***||^2^*_Fro_*, yielding 73.5%.

For the NMF analysis of axonal volumes shown in **Figure 3**, axon voxels outside the region of interest (ROI), such as the CP or GPe, were excluded to prevent non-ROI signal from influencing the analysis. ROI voxels were smoothed using a 3D Gaussian filter (σ = 100 μm) and down-sampled to 25-μm voxel resolution. The smoothed volumes were linearized and concatenated into a 2-D matrix (e.g., 17 brain samples × 906,439 CP voxels for the CP analysis). This matrix was then analyzed using the same NMF procedure described above, with a randomly selected 10% of voxels from the Gaussian-filtered ROI volumes designated as the test set and the remaining voxels used for training. We note that RMSE_Replicate_ was not computed for these voxel-level analyses, as the injection-site-label-based optimal model lacks predictive power at the voxel level and yields error indistinguishable from RMSE_Shuffle_. The resulting basis pattern matrices were reconstructed in 3-D for biological interpretation. For clustering analyses based on scores from NMF decomposition, the optimal number of clusters was determined using the gap statistic^112^ for hierarchical clustering unless otherwise specified. To evaluate the robustness of the clustering shown in **Figure 2**, we performed additional clustering analyses using alternative methods (**Figure S2**). The optimal number of clusters for hierarchical clustering, *k*-means clustering, and Gaussian mixture models was determined using the MATLAB function “evalclusters.” For spectral clustering, which is not supported by “evalclusters,” the optimal cluster number was determined using the “silhouette” function.

#### Marker region analysis

To identify marker regions across the 280 brain regions (**Figure S4**), we first grouped the 110 brain samples into their respective projectomic groups. Regional axon data were normalized by the total axon quantity within each brain (analogous to "counts-per-million" in single-cell RNA sequencing). For each region, we performed a one-way ANOVA across the five projectomic groups and identified candidate "sole-cluster" regions—those in which the highest group mean was at least double the means of all other four groups. We then applied the two-stage Benjamini-Krieger-Yekutieli (BKY) procedure^113^ to control the false discovery rate at q < 0.05 across all 280 regions, retaining only those candidates whose ANOVA q-values passed this threshold. Finally, to confirm selective enrichment, we performed Welch’s two-sided t-tests comparing the sole-cluster group against each of the other four groups (four comparisons per region), again corrected by the BKY procedure within each region; only regions in which all four post-hoc q-values were below 0.05 were retained as marker regions. Using this procedure, we identified 16, 8, 18, 28, and 81 marker regions for the HPC, BG, CTX, MI, and BSLT groups, respectively.

#### Supervised NMF analysis

For the supervised NMF analysis of projection-specific distributions of labeled cell bodies within the raphe nuclei, we first binned the cell distribution data to a voxel size of 25 μm × 25 μm × 25 μm. We noted that the DR boundary defined by the CCFv3 was somewhat too narrow in the coronal planes. Therefore, we included the labelled cells in the ventrolateral portion of the periaqueductal gray as DR in our analysis. The data were smoothed using a 3-D Gaussian filter (σ = 125 μm) and organized into a 2-D matrix in the same manner as the axonal volume analysis. To further save computational resources, we removed voxels that were empty across all samples. This process resulted in a 2-D matrix of size 110 samples × 983,686 voxels.

The main NMF algorithm remained largely the same with a few modifications. First, the cost function of the original NMF algorithm^35,36^ was adjusted to incorporate classification:

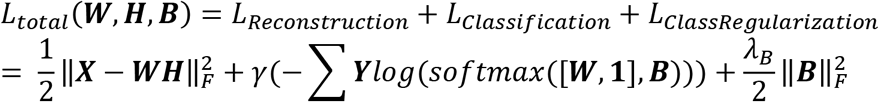

where ***X****, **W**,* and ***H*** represent the smoothed cell body distribution matrix, score matrix, and basis matrix, respectively; ***Y*** and ***B*** denote the one-hot encoded label matrix and the classification matrix, respectively; and *γ* and λ are the regularization parameters for classification and ***B***, respectively. Note that we introduced a bias vector to incorporate the bias term for classification. Our aim was to solve the following equation:

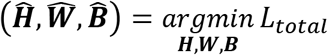

The next modification we made is that, while the three matrices, ***W***, ***H***, and ***B***, were randomly initialized with non-negative values, these matrices were iteratively updated according to the following update rules, in which the classification components were derived by taking the derivative of the loss function with respect to the relevant matrices:

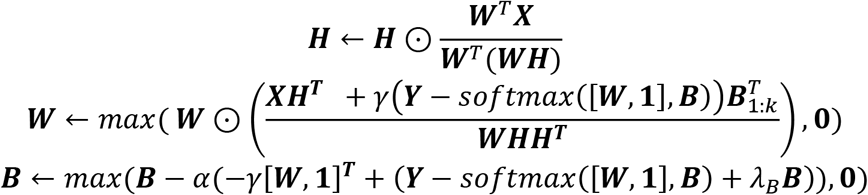

where ⊙ denotes element-wise multiplication; *k* denotes the number of bases (i.e. latent factors); *n* and *m* denote the sample- and original feature dimensions, respectively; *α* denotes the learning rate for gradient descent. Note that the update rule for ***H*** and the first term in the ***W*** update rule, ***XH^T^***, are derived from the original update rule^36^.

To find the optimal set of hyperparameters, we employed a 5-fold cross-validation and Bayesian optimization. Using the obtained set of hyperparameters, **Figure S7B** and **C** were generated using the proposed supervised NMF algorithm on all 110 samples without cross-validation.

#### STARmap in situ sequencing

STARmap was performed as previously described^68,114^. Briefly, animals were injected with AAV_8retro_ encoding circular mRNA barcodes^101^ at multiple sites in the right hemisphere (packaged by Neurotools; injection regions, virus titer, and volumes are listed in **Table S4**). In total, 14 B6 wild-type mice (one male and one female per injection combination) received retrograde barcode injections into 24 brain regions spanning the five projectomic groups (**Table S4**). To enable both between- and within-projectomic-group analyses, mice were divided into two groups: four “cross-group” mice received injections at brain regions of all five projectomic groups (serving as a baseline for cross-group co-projection), and each of the ten “within-group” mice (two mice per projectomic group) received injections restricted to brain regions of a single projectomic group. Three to four weeks after injections, animals were euthanized using carbon dioxide followed by cervical dislocation. Brains were dissected and embedded in O.C.T. (23-730-571, Fisher Scientific), then snap-frozen in liquid nitrogen vapor and stored at −80°C until use.

Tissue blocks were sectioned into 10-µm slices mounted onto silanized well plates (P24G-1.5-13-F, MatTek), fixed with 4% PFA for 10 minutes, washed in 1× PBS, and permeabilized with methanol for 1 hour at −80°C. Sections were rehydrated in PBSTR (0.1 U/µL SUPERasin; 1× PBS; 0.1% Tween 20), hybridized overnight at 40°C (0.01 µM each probe, **Table S3**; 2× SSC; 10% deionized formamide; 1% Tween 20; 20 mM RVC; 0.1 mg/mL salmon sperm DNA), and processed through ligation (T4 DNA ligase, 2 hours) and rolling circle amplification (Phi29 DNA polymerase, 4 hours at 30°C). Amplicons were embedded in a polyacrylamide gel (4% acrylamide, 0.2% bis-acrylamide), and tissue was digested with Proteinase K (0.2 mg/mL, 1 hour at 37°C).

We designed padlock probes targeting two serotonergic markers (*Tph2* and *Slc6a4*), 10 transcriptomic subtype marker genes (*Syt10*, *Sox14*, *Gad1*, *Syt2*, *Npas1*, *Tacr3*, *Ppp1r17*, *Irx2*, *Met*, and *Piezo2*; probe sequences in **Table S3**), and five retrograde circular barcodes. For each of six sequencing rounds, sections were incubated in ligation mix (T4 DNA ligase, reading probe, and fluorescent oligos) for 3 hours, washed in imaging buffer (2× SSC, 10% formamide), and imaged using a Nikon Eclipse spinning disk microscope with a 40× oil lens (Plan Fluor 40×/1.30 oil, WD 0.24). Fluorescent signals were stripped between rounds with 80% formamide and 0.1% Triton-X-100.

#### STARmap data processing

Preprocessing was performed in MATLAB 2021b. After sequencing, 3D image stacks comprising four channels (DAPI at 405 nm and three signal channels at 488, 546, and 647 nm) across six imaging rounds were restitched by re-computing optimal tile overlap using Fourier-transform-based cross-correlation. Amplicon positions were extracted using a Laplacian-of-Gaussian filter with manually set channel thresholds, and spot-calling results were inspected to confirm the absence of between-round or cross-channel bleed-through. Rounds were aligned using the DAPI channel with both linear and non-linear registration, and transformations were applied to all channels. Cell segmentation was performed using Cellpose^105^, and amplicons were assigned to cells using Baysor^106^, generating a cell-by-gene count matrix.

Serotonin neurons were identified based on *Tph2* expression exceeding a raw count threshold of 20. Cells with fewer than 3 total counts across the 10 subtype marker genes were excluded from the downstream analyses. This initial screening yielded 8,715 serotonin neurons across 14 mice. Of these, 635 cells (7.3%) were barcode-positive for at least one target, distributed across groups as follows: 227 cross-group, 65 HPC, 61 BG, 35 CTX, 177 MI, and 70 BSLT.

Gene expression counts for the 10 marker genes were log-transformed (log(1 + x)) and z-scored across all cells. To correct for inter-animal batch effects, Harmony^108^ was applied using mouse identity as the batch variable (maximum 25 iterations). Principal component analysis was performed on the Harmony-corrected expression matrix, retaining components explaining >90% of cumulative variance (9 PCs). Unsupervised clustering was performed using the Leiden algorithm on a k-nearest-neighbor graph (k = 15, correlation distance) constructed from the PCA-reduced data.

Serotonin neurons were mapped to 10 previously defined transcriptomic subtypes^25^ (cluster 4 excluded due to absence in a preliminary whole-brain imaging experiment using *Sert-Flp;Crh-Cre;Ai65*) using a two-stage consensus classification strategy. In the first stage, each cell was assigned to one of six mega-types (groups of related subtypes: *Gad1* group [Clusters 1, 2, 3]; *Syt2/Npas1* group [Clusters 5, 6]; *Tacr3* group [Clusters 7, 10]; and singletons *Ppp1r17* [Cluster 8], *Irx2* [Cluster 9], *Met/Piezo2* [Cluster 11]) using four independent methods: 1. KNN classification (k = 5, correlation distance) trained on z-scored reference data; 2. Centroid-based assignment using correlation distance to mega-type centroids computed from the reference dataset; 3. Leiden cluster transfer, in which each STARmap Leiden cluster was mapped to a reference mega-type by correlation of cluster-mean expression profiles, and cells inherited the assignment of their cluster; 4. Quantile-normalized KNN, in which STARmap gene expression was quantile-normalized to the reference distribution prior to KNN classification. A cell was assigned to a mega-type only when at least three of four methods agreed. In the second stage, cells within multi-subtype mega-types were further resolved to fine subtypes using the same four-method consensus applied within each mega-type, with method-specific models retrained on the relevant reference subset. Cells failing to reach consensus at either stage were excluded. This procedure yielded 6,214 high-confidence cells (71.3%). To evaluate classification performance, the two-stage strategy was applied in 5-fold cross-validation on the reference dataset^25^ (775 cells). The strategy achieved 75.9% accuracy among high-confidence predictions, with 93.3% of cells meeting the confidence threshold.

To identify barcode-positive cells, we applied a data-driven thresholding approach. For each mouse and barcode channel, we swept a range of raw count thresholds. At each threshold, we computed Cramér’s V—the association strength between barcode-positive status and transcriptomic subtype composition—by comparing the subtype distribution of thresholded barcode-positive cells to the baseline distribution of all confident cells via a chi-square statistic. The threshold maximizing Cramér’s V (requiring a minimum of 5 barcode-positive cells) was selected. This approach adaptively sets thresholds per mouse and channel, accounting for variation in viral expression and injection efficiency.

To test whether specific transcriptomic subtypes were preferentially associated with particular projectomic groups, we computed the enrichment ratio (ER) for each subtype–group pair:

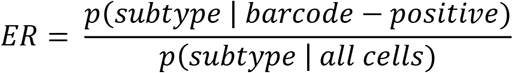

where ER > 1 indicates over-representation and ER < 1 indicates under-representation. Statistical significance was assessed using Fisher’s exact test on the 2 × 2 contingency table (subtype membership × barcode-positive status). Cells from all mice contributing barcode-positive cells for the relevant group were pooled. Subtype–group pairs with fewer than 5 cells in either the subtype or barcode-positive category were excluded (approximately 10 tests per group). P-values were corrected using the two-stage Benjamini-Krieger-Yekutieli (BKY) false discovery rate procedure^113^ (p = 0.05), applied independently within each projectomic group. Replicate consistency was assessed by computing the odds ratio separately for each cohort (one male, one female); an enrichment was considered replicate-consistent if the odds ratio exceeded 1 in both cohorts.

To assess within-group axonal collateralization, we compared the multi-barcode rate—the proportion of barcode-positive cells carrying two or more barcode types—between within-group mice (injected at multiple sites belonging to the same projectomic group) and cross-group mice (injected across different groups). For each within-group projectomic group, the multi-barcode rate was compared to the cross-group baseline using a difference permutation test (50,000 iterations). In each permutation, barcode channel labels were simultaneously shuffled for both within-group and cross-group mice, preserving the channel configuration and mouse structure of the original experiment. The observed difference in multi-barcode rate (within-group minus cross-group) was compared to the null distribution to obtain a one-sided p-value. P-values were corrected across the five projectomic groups using BKY FDR (p = 0.05). Error bars in co-projection figures represent the binomial standard error of the observed proportion.

Beyond transcriptomic and barcode-based analyses, we also extracted spatial information for each cell. Each brain section was imaged at 10× magnification to capture the entire view of the section. To obtain 3-D coordinates of analyzed cells in a reference space, we first linearly registered the high-magnification samples to the 10× overview images and then used AMaSiNe^107^ to localize these sections within the Allen CCFv3^34^. We trained a random forest classifier to predict the projectomic group of individual neurons from their spatial coordinates, transcriptomic cluster identity, or both (**Figure 5H**; **Figure S8G, H**). Cells whose barcodes mapped to more than one projectomic group were excluded from classifier training and evaluation, yielding n = 430 single-group-labeled cells.

UMAP dimensionality reduction was computed using densMAP on the PCA-reduced expression matrix of confident cells. Harmony, Leiden clustering, and UMAP were implemented in Python (harmonypy, scanpy, umap-learn). All statistical analyses and figure generation were performed in MATLAB.

#### Intersectional validation of Met/Piezo2 neurons

To independently validate the association between the Cluster 11 subtype and specific projectomic groups, *Piezo2-Cre;Sert-Flp* double-transgenic mice (6–8 weeks old) were injected with a Cre/Flp-dependent mCherry reporter (AAV-Con/Fon-mCherry; 137132-AAV8, Addgene) into the raphe. Six weeks after injection, mice were sacrificed and brain sections were stained with antibodies against RFP (1:1000, 600-401-379, Rockland) and Tph2 (1:500, ab121013, Abcam). Labeled neurons were assessed for co-expression with Tph2, and projection targets were identified by visualizing mCherry-positive axons in serial brain sections.

#### Animal behavior

*Tph2^fl/fl^;Sert-Flp;Ai65* mice were used as the experimental group and *Sert-Flp;Ai65* mice served as controls. Both male and female mice aged 6–8 weeks underwent stereotaxic viral injection. Mice were single-housed 14 days after surgery, and behavioral testing started 3 weeks post-injection. Mice were maintained on a 12:12 h light/dark cycle (lights on at 7 AM). All behavioral assays were conducted during the light phase (9 AM-5 PM) at approximately 100 lux unless otherwise noted except the open-field test, which was performed during the dark phase (8 PM-6AM) under red light illumination.

#### Surgical procedures for behavior experiments

Mice underwent stereotaxic viral injection as described above. A custom AAV_8retro_ construct was generated by replacing the EF1a promoter in pAAV-EF1a-FLEX_FRT_-Cre (121675, Addgene) with the CAG promoter and packaging into the AAV_8retro_ capsid (titer 2.4e13). For Batch 1 samples, virus was bilaterally injected into one of five target regions: dorsal CA3 (dCA3; HPC group), globus pallidus externus (GPe; BG group), piriform cortex (PIR; CTX group), lateral preoptic area (LPO; MI group), or superior colliculus (SC; BSLT group). Control mice received injections distributed across all five regions (one male and one female per region). Sample sizes (male, female) were as follows: dCA3 (5, 4), GPe (6, 4), PIR (4, 6), LPO (5, 5), SC (5, 5), and control (5, 5).

For Batch 2, virus was injected into a different set of target regions within the same broader anatomical categories: ventral CA1 (vCA1; HPC group), substantia nigra pars reticulata (SNr; BG group), agranular insular cortex (AI; CTX group), paraventricular thalamus (PVT; MI group), or lateral geniculate nucleus (LG; BSLT group). Sample sizes for Batch 2 (male, female) were: vCA1 (2, 2), SNr (3, 1), AI (3, 1), PVT (1, 3), LG (2, 2), and control (3, 3). Control mice were distributed across regions, with males injected in vCA1, SNr, and AI, and females in vCA1, AI, and LG. Injection coordinates and volumes for all target regions are listed in **Table S1**.

#### Marble burying test (MBT)

The marble burying test was performed following the adapted protocol previously described^76^. Mice were individually placed in plastic disposable cages (L30 cm × W18 cm × H12.5 cm) containing fresh, unscented bedding at a depth of 3.5 cm. Twenty blue glass marbles (16 mm diameter) were evenly arranged in a 4 × 5 grid on the leveled bedding surface. Prior to testing, mice were habituated to the testing room in their home cages for 30 minutes under identical lighting conditions. Each mouse was then gently transferred to the experimental cage, which was covered with a transparent acrylic lid to minimize airflow disturbance while allowing visual monitoring. Behavior was recorded for 30 minutes using an overhead video camera. Bedding was used only once, and marbles were washed with 70% ethanol and completely dried. At the end of the session, the number of marbles buried (defined as ≥ 2/3 of the marble surface covered by bedding) was scored by observers blinded to experimental group.

#### Elevated plus maze (EPM)

The elevated plus maze (EPM) consisted of two open arms and two closed arms (30 × 5 cm each) extending from a central platform, elevated 85 cm above the floor. The maze was illuminated at 200 lux. At the start of each trial, the mouse was placed at the distal end of an open arm facing the center and allowed to freely explore for 5 minutes^82^. Between subjects, the maze surface was cleaned with 70% ethanol and allowed to dry completely before the next trial.

#### Open field test (OFT)

The behavioral apparatus and acquisition setup were constructed following the specifications described on the MoSeq2 hardware wiki (https://github.com/dattalab/moseq2-app/wiki), and the experimental protocol was adapted from a previous study with a modification in session duration^79^. An overhead depth camera (Microsoft Kinect) was frame-synchronized with a centrally positioned collimated LED (4900 K, 740 mW; MNWHL4 and SM2F32-A, ThorLabs) driven by a DC4104 driver (ThorLabs) and controlled via MicroManager software.

All experiments were conducted during the dark phase (8 PM-6 AM) in a room maintained under red light with continuous white noise. Each mouse was placed in the center of a circular open-field arena (catalog no. 14317, US Plastic) using a 5-inch net pot to minimize handling stress and bedding transfer. Mice were allowed to move freely throughout the arena. Video acquisition began immediately upon placement. Each session consisted of four 10-minute phases: an early light-off period, a late light-off period, a center light-on period in which the LED illuminated approximately 40% of the total arena area (∼65% of the radius from center), and a final re-light-off period. Between subjects, the arena was cleaned sequentially with 10% Alconox (catalog no. 21835-032, VWR), water, and 70% ethanol, and allowed to dry completely before the next trial. Depth recordings were preprocessed using the MoSeq2 pipeline^88^, and scalar summary statistics were used for subsequent analyses.

#### Sucrose preference test (SPT)

The two-lickometer sucrose preference test apparatus consisted of a Bpod State Machine r2 (product ID 1024, Sanworks) and two infrared-based lick ports (product ID 1004, Sanworks) mounted in a plastic cage (L30 cm × W18 cm × H12.5 cm). The two lickometer ports were mounted on the same side of the cage, spaced 8 cm apart and positioned 3 cm above the floor. The cage was covered with a transparent acrylic lid to prevent escape.

Mice were water-deprived for 16 hours with food available ad libitum prior to the habituation session. During habituation, mice were placed in the apparatus and given free access to plain water and sucrose water (2% w/v) from the two ports for two hours (3 μl dispensed per bout). Port assignment (plain vs. sucrose) was randomized across mice but kept consistent between the habituation and test sessions for each mouse. Following habituation, mice were returned to their home cages with ad libitum access to food and water for one day of recovery. Mice were then water-deprived again for 24 hours with ad libitum food access, after which they underwent the same two-hour two-lickometer choice test. Between subjects, the lickometer ports were rinsed with water and the apparatus was cleaned with 70% ethanol, and both were allowed to dry completely before the next mouse. Lick timestamps and port identity were recorded for subsequent analysis.

#### Feeding test

Feeding behavior was assessed using an automated pellet dispenser (FED3.2, Open Ephys) loaded with 20 mg dustless precision pellets (catalog no. FF-0071, Bio-Serv). The device dispensed a single pellet each time the mouse performed a nose poke at the pellet retrieval port.

The experiment consisted of three phases. During a 3-day acclimation phase, standard chow in the home cage was replaced with 20 mg pellets to familiarize mice with the novel food. This was followed by a training session in which mice were food-deprived for 24 hours and then placed in the experimental setup with the FED3 device for 2 hours to acquire the nose-poke response required for pellet retrieval. After training, mice were returned to their home cages with food and water available ad libitum for 24 hours of recovery. For the test session, mice were again food-deprived for 24 hours and feeding behavior was recorded over a 2-hour session using the FED3 device. Between subjects, the nose-poke port and pellet retrieval well were rinsed with water, and the apparatus was cleaned with 70% ethanol and allowed to dry completely before the next mouse. Pellet dispense timestamps were logged by the FED3 device for subsequent analysis.

#### Auditory fear conditioning (FC)

The fear conditioning protocol was adapted from a previous study^24^. The conditioning chamber consisted of a square enclosure (L18 × W18 × H30 cm) with a metal grid floor connected to a shock generator and scrambler, housed within a sound-attenuating chamber (Coulbourn Instruments). To create distinct contexts, the transparent chamber walls were lined with one of two wallpaper patterns, and one of two odorants (peppermint oil, W284807; or artificial garlic oil, W530316; both Sigma-Aldrich), diluted 1:2000 in mineral oil, was placed below the grids in the chamber. The wallpaper–odor combination was randomly assigned across mice, with the same combination used for conditioning and contextual recall, and a different combination used for cue recall. An auditory tone (6 kHz, ∼75 dB, 30 s) served as the conditioned stimulus (CS).

On day 1 (conditioning), mice were placed in the chamber and allowed a 3-minute baseline period to habituate to the context. Mice then received 3 CS presentations (30 s each, 60 s inter-trial interval), each co-terminating with a 1-s foot shock (0.5 mA). Mice remained in the chamber for an additional 60 seconds after the final CS–US pairing. On day 2, contextual fear memory was assessed by returning mice to the conditioning chamber with the original wallpaper and odor for 5 minutes in the absence of any auditory cues. Two hours later, cue-dependent fear memory was tested by placing mice in the chamber with a novel wallpaper–odor combination. Following a 3-minute baseline exploration period, 8 CS presentations were delivered without foot shocks (15 s duration, 30 s inter-stimulus interval). Between subjects, the chamber walls, grid floor, and waste tray were cleaned with 70% ethanol and allowed to dry completely before the next mouse.

#### Tail suspension test (TST)

The tail suspension test (TST) was performed to assess coping strategy following the modified protocol from a previous study^115^. Mice were individually suspended by the tail from a horizontal bar positioned approximately 35 cm above the surface using adhesive tape applied approximately 1 cm from the tip of the tail. Each session lasted 6 minutes and was recorded by a side-mounted video camera. Between subjects, the adhesive tape was replaced and the apparatus was cleaned with 70% ethanol and allowed to dry completely before the next mouse.

#### Forced swim test (FST)

The forced swim test (FST) was conducted over two consecutive days following the protocol described in a previous study^24^. Mice were individually placed in a plastic cylinder (25 cm height, 18 cm diameter) filled with water (23 ± 1°C) to a depth of 15 cm, ensuring that mice could not support themselves by touching the bottom with their hind limbs. On day 1, mice underwent a pre-exposure swim session. On day 2, mice were tested under identical conditions and behavior was recorded for 6 minutes. After each session, mice were gently dried with absorbent paper towels and placed on a heating pad until fully dry before being returned to their home cages. Between subjects, water was replaced and the cylinder was rinsed to remove residual odor cues.

#### Behavioral video processing

Video recordings from the MBT, EPM, FC, TST, and FST were analyzed using a custom deep-learning-based pipeline that operates directly on raw video frames to reliably detect subtle behaviors such as grooming in the MBT, head dipping in the EPM, and immobility in the FST. Behavior was classified using a two-stage deep learning pipeline. In the first stage, a YOLOv8 object detection model (Ultralytics) was trained to detect and localize the mouse in each video frame. A fixed-size crop (224 × 224 pixels) was extracted around the center of the detected bounding box, and when detection failed on a given frame, the bounding box from the most recent successful detection was carried forward to maintain spatial continuity. Training frames for the YOLO model were obtained by randomly sampling frames from experimental videos and manually annotating them.

For behavior classification, behavioral segments were independently annotated by two observers using a custom annotation tool, and the resulting JSON annotation files were used to extract short video clips (annotation tool at https://github.com/tshindmarsh/TUBBA). Clips were sorted into behavior-specific subdirectories to form the training and validation sets. Training clips were generated using a sliding window with a configurable stride to allow temporal overlap, whereas validation clips were extracted without overlap. Training and validation sets were split at the level of individual animals (by mouse ID) to prevent data leakage.

The behavior classifier was a SlowFast network^89^ pretrained on Kinetics-400. The original classification head was replaced with a three-layer fully connected head (512–256–N classes, with ReLU activations and dropout of 0.5) and fine-tuned using a progressive unfreezing strategy, in which deeper backbone blocks were gradually unlocked across successive training stages. Training employed data augmentation (random horizontal flip, affine transformations, color jitter, Gaussian blur, random erasing, and additive/multiplicative noise) and the AdamW optimizer with differential learning rates. The automated behavioral classifiers were validated against manual scoring, yielding macro-averaged F1 scores of 0.86 for the marble burying test (four classes: marble burying, grooming, rearing, and other), 0.87 for the elevated plus maze (head dipping vs. other), 0.86 for fear conditioning (freezing vs. non-freezing), 0.87 for the tail suspension test (active coping vs. immobility), and 0.96 for the forced swim test (active coping vs. immobility).

At inference, YOLO detection and SlowFast classification were applied sequentially to each full-length video. The SlowFast model received fixed-length clips via a sliding window and assigned a behavior label to the center frame of each clip based on softmax probabilities. A post-processing filter removed events shorter than a minimum duration threshold to eliminate implausibly brief behavioral bouts. The pipeline output consisted of frame-by-frame behavior labels, class probabilities, and bounding box coordinates, saved as CSV files.

All analysis code was written in Python using PyTorch, PyTorchVideo, Ultralytics YOLOv8, and OpenCV, and is available on GitHub https://github.com/junhsong92/serotonin_projectome.

#### Behavioral feature definition and selection

The 54-feature set (**Figure S9A**, left) comprised, by paradigm: the marble burying test (4 features: total number of marbles buried, digging duration, grooming duration, and rearing duration); the open field test (8 features: thigmotaxis duration and average movement speed, each measured separately across the four protocol phases — early light-off, late light-off, center light-on, and final re-light-off); the elevated plus maze (4 features: total distance moved, head dip count, open-arm entrance count, and open-arm duration); the sucrose preference test (13 features: total licks; sucrose preference ratio computed over the whole 2-hour session; fraction of sucrose licks during the first 30 minutes relative to total sucrose licks; fraction of sucrose licks during the last 30 minutes relative to total sucrose licks; fraction of water licks during the first 30 minutes relative to total water licks; fraction of water licks during the last 30 minutes relative to total water licks; fraction of total liquid licks during the first 30 minutes; fraction of total liquid licks during the last 30 minutes; sucrose preference ratio during the first 30 minutes; sucrose preference ratio during the last 30 minutes; preference change index, defined as the difference between first 30-minute and last 30-minute preference ratios; time required to reach 50% of total sucrose licks; and time required to reach 50% of total water licks); the feeding test (9 features: body-weight-normalized total food consumption, average inter-feeding interval, latency to first feeding bout, average pellet consumption rate, ratio of pellets consumed in the first 30 minutes, ratio of pellets consumed in the last 30 minutes, satiety index (calculated as [P_early − P_late]/[P_early + P_late], where P_early and P_late are the number of pellets consumed during the first and last 30 minutes of the session), time required to reach 50% of total food consumption, and coefficient of variation of inter-feeding intervals); fear conditioning (9 features: freezing during the last session of conditioning; freezing during the first half of contextual recall; freezing during the second half of contextual recall; difference in freezing between the first and second halves of contextual recall; freezing across the whole contextual recall session; freezing during the pre-onset baseline of cued recall; freezing averaged across the first two cued recall trials; freezing averaged across the last two cued recall trials; and difference in freezing between the first and last cued recall trials); the tail suspension test (2 features: active coping duration and the temporal area under the curve of active coping); and the forced swim test (5 features: inactive duration on Day 1, inactive duration on Day 2, difference in inactive duration between Day 2 and Day 1, temporal area under the curve of inactive duration on Day 1, and temporal area under the curve of inactive duration on Day 2).

Features were defined a priori based on the standard outputs of each behavioral paradigm and its corresponding scoring pipeline (MoSeq scalar outputs for the open field test, SlowFast classifier outputs for the marble burying test, the elevated plus maze, fear conditioning, the tail suspension test, and the forced swim test; FED3 logs for the feeding test; and lickometer recordings for the sucrose preference test). No feature was added or removed after observing group differences in the data, avoiding feature-selection-based circularity.

Within each paradigm, we included readouts that captured three aspects of behavior. The first is the overall amount or rate of a behavior across the entire session, such as the total number of marbles buried in the marble burying test or the overall sucrose preference ratio in the sucrose preference test. The second is how a behavior is distributed within a single session, such as the fraction of sucrose licks during the first 30 minutes versus the last 30 minutes of the sucrose preference test, or the temporal area under the curve of active coping and inactive duration in the tail suspension test and the forced swim test. The third is how a behavior changes across separate sessions, such as the difference in inactive duration between Day 2 and Day 1 of the forced swim test, or the difference in freezing between the first and second halves of contextual recall in fear conditioning. Including these alongside single-summary metrics allowed us to detect group differences not only in the overall magnitude of a behavior but also in when it occurs within a session and how it changes over repeated sessions.

#### Multivariate behavioral classification using Linear Discriminant Analysis

To identify a combination of behavioral features that best discriminates between experimental groups, we developed a supervised classification pipeline in MATLAB using Linear Discriminant Analysis (LDA) with leave-one-out cross-validation (LOOCV). Given the sample size relative to the dimensionality of the behavioral feature set, multiple safeguards were implemented to prevent overfitting throughout the pipeline.

Behavioral features were z-scored within each sex independently to account for sex-specific differences in baseline behavior. Z-scored values exceeding ± 5 standard deviations were winsorized to the cap value. Missing values were sparse and limited to edge cases such as undefined inter-event intervals when only a single pellet was consumed during the feeding experiment; these were imputed using k-nearest neighbors (k = 3).

Feature filtering for the LDA classifier was performed in two stages. As a first step toward overfitting prevention, features were individually tested for group differences using a bootstrap ANOVA (50,000 permutations), and only features reaching significance (p < 0.05) were retained. The surviving features were then ranked by their contribution to between-group separation, quantified as the weighted sum of absolute LDA loadings scaled by the corresponding eigenvalues, with L2 regularization applied to the within-class scatter matrix to further constrain the model.

The optimal number of features and discriminant dimensions was determined via an exhaustive grid search over all combinations of ranked feature counts (up to a maximum of 10, given the sample size) and LDA dimensions (up to k − 1, where k is the number of groups), selecting the combination that maximized overall LOOCV accuracy. Following this initial selection, an iterative feature injection procedure was applied to improve the minimum per-class accuracy. This criterion was chosen because overall accuracy can mask poor classification of individual groups; optimizing for the worst-performing class ensures that the model does not achieve high aggregate accuracy at the expense of failing to distinguish specific experimental conditions. At each iteration, the most confused class pair was identified from the confusion matrix, and candidate features from the remaining pool were ranked by Cohen’s d for that pair. The candidate yielding the greatest improvement in minimum class accuracy was added to the feature set. This process continued until no further improvement was found or the feature limit was reached.

Classification performance was assessed using LOOCV with nearest-centroid classification in the LDA-projected space. To assess statistical significance, a permutation test (N = 10,000) was conducted in which group labels were shuffled and the full pipeline, including grid search and feature injection, was re-run for each permutation. This shuffled-label control provided an empirical null distribution against which observed accuracy could be tested, addressing the possibility that the multi-step feature selection itself inflates classification performance. The p-value was computed as the proportion of permutation accuracies meeting or exceeding the observed accuracy. Notably, the resulting empirical chance level (mean of the permutation distribution: ∼26%) exceeds the naive theoretical chance of 20% for a 5-way classification. This difference reflects the contribution of the multi-step feature selection itself: when applied to shuffled-label data, the pipeline can still capture a small amount of apparent structure due to data-driven feature ranking and injection. By using the empirical permutation null rather than 20%, we conservatively account for this pipeline-induced baseline inflation when evaluating significance.

For independent replication (Batch 2), behavioral features were z-scored independently using Batch 2 statistics. Covariance alignment (CORAL^93^) was then applied to transform Batch 2 data to match the covariance structure of Batch 1 before projecting onto the discriminant axes derived from Batch 1.

## SUPPLEMENTAL VIDEO AND EXCEL TABLE TITLES AND LEGENDS

**Video S1. Fly-through videos showing whole-brain axon innervation patterns of two sample brains, related to Figure 1**

Videos showing serial coronal sections from anterior to posterior for the two brains shown in Fig. 1C. Injection sites at dorsal striatum (left) and dorsal hippocampus (right) are indicated by *. CCFv3 (10-μm resolution) was upsampled to a 5-μm resolution and cropped to approximately match our imaging window. Then, 1,890 coronal virtual sections, each 5 μm thick, were overlaid with the processed axon volumes for visualization.

**Table S1. Injection site coordinates and injection volume for axon tracing and behavior experiments, related to Figures 1, 6, and 7**

Injection site coordinates and injection volume used to initiate axon tracing are provided. We note that the five injection sites in the CP were selected without prior knowledge of the results described in Fig. 3 but were chosen to ensure they were evenly distributed throughout the CP.

**Table S2. Raw data for axon density and volume for 110 sample brains across 280 brain regions, related to Figures 1 and 2**

Raw data and L2-normalized axon density and quantity are shown in separate sheets. Brain region abbreviations are given in Column B. Brain IDs are given in parentheses in Row 1.

**Table S3. Probe sequences for STARmap, related to Figure 5**

Probe sequences for marker genes and barcodes (Tab 1) and orthogonal reading (OR) probes (Tab 2) for spatial transcriptomic experiments using STARmap are provided. The sequences for *Gad1* were adapted from a previous study^114^.

**Table S4. Detailed information about retrograde virus encoding circular barcodes, related to Figure 5** Information regarding retrograde viruses encoding circular barcode sequences, injection region combinations, and injection volumes is provided, with injection coordinates detailed in Table S1.

**Table S5. STARmap expression, related to Figure 5**

Retrograde barcode and marker gene puncta counts, individual cell coordinates, and the mouse and slice IDs of the cells used in the STARmap analyses are provided.

**Table S6. Behavior quantification, related to Figures 6 and 7**

Quantified behavioral features for all mice across the eight behavioral paradigms are provided. Mouse IDs, projectomic group assignments, sex, batch, and control/experimental designations are included.

---

**Figure S1.**
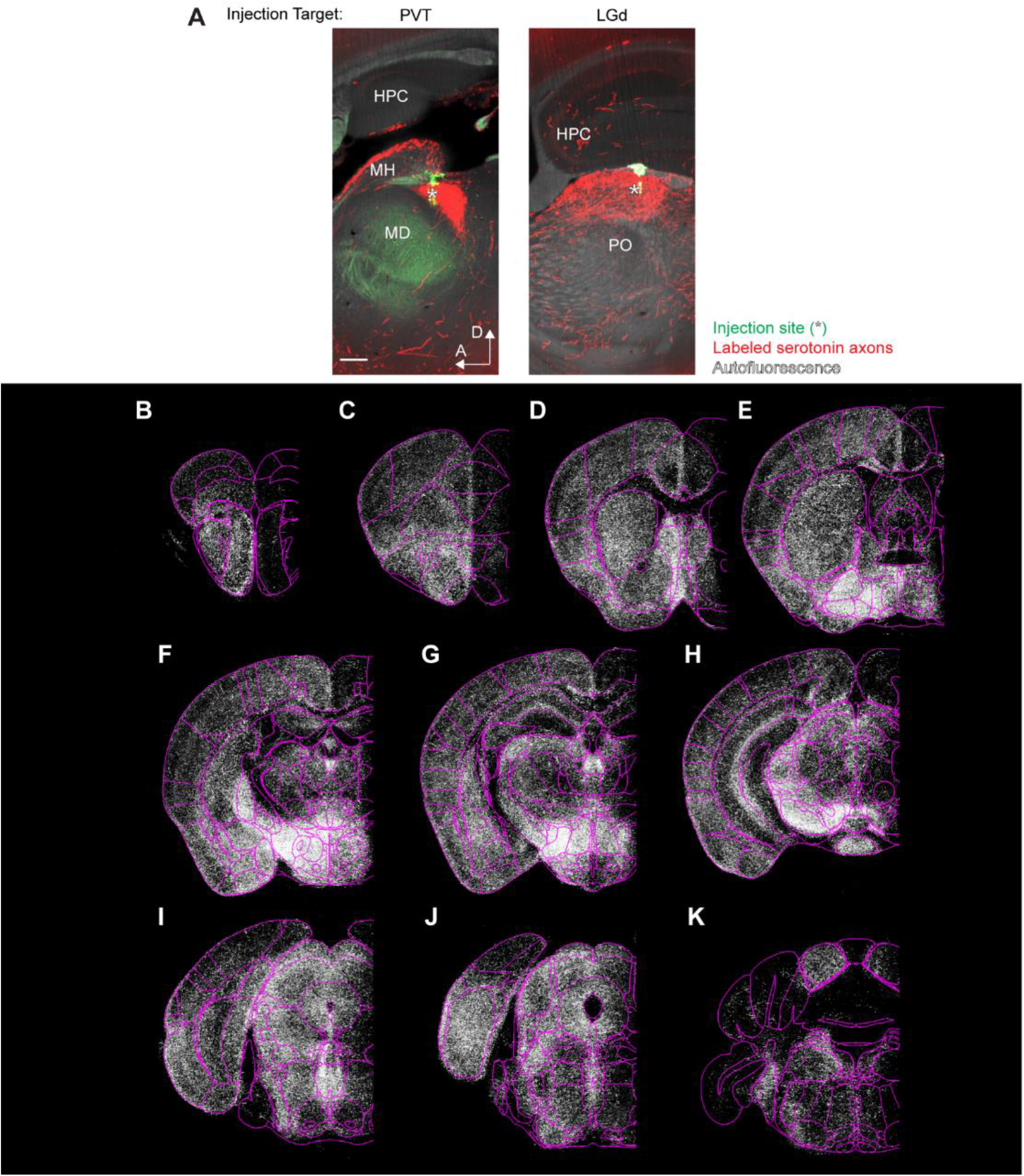
Validation of injection specificity and whole-brain axon coverage of the dataset, related to Figure 1. **(A)** Potential virus leakage and spillover into brain regions adjacent to the intended injection sites were examined. For example, the mediodorsal nucleus of the thalamus (MD), a region neighboring the paraventricular nucleus of the thalamus (PVT), shows little axon labeling in samples where the virus was injected into the PVT (left panel). Similarly, the posterior complex of the thalamus (PO) exhibits no dense labeling in samples injected into the dorsal part of the lateral geniculate complex (LGd; right panel). These data confirm that the viral injections remained mostly confined to their designated targets with minimal unintended transduction in neighboring brain regions. Red retrobeads, pseudocolored green, were mixed with the virus at a 1:25 ratio to mark the injection sites. Scale bar: 200 µm; Section thickness: 30 µm; maximal intensity projection. HPC, hippocampus; MH, medial habenula. **(B–K)** Overlay of axons segmented from all 110 sample brains in the dataset (white). Example coronal slices were collected along the anterior-posterior (AP) axis at 300–500 µm intervals. Magenta outlines denote the boundaries of brain regions used in analyses. Note that the contralateral hemisphere (right) was not fully imaged due to the working depth limitations of the objective used.

**Figure S2.**
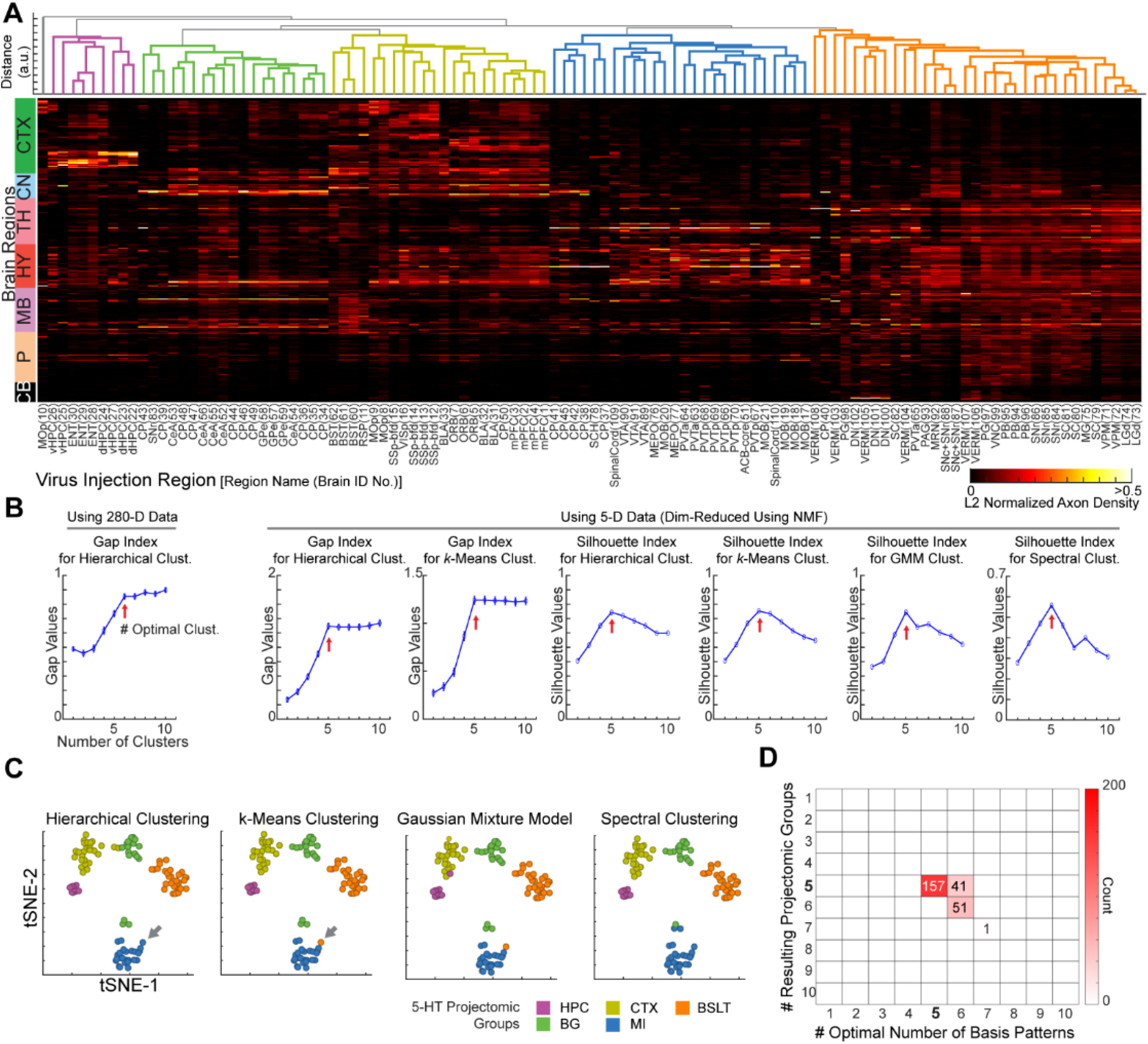
Evaluation of the stability of the NMF-based analysis, related to Figure 2. **(A)** Hierarchical clustering using the 280-D axon density data shown in Figure 1D. Hierarchical clustering was performed on samples in the original dimensional space (110 brains) to evaluate whether the clustering results shown in Figure 2E, based on NMF-dimension-reduced samples, accurately reflect clustering using 280-D normalized axon density data. Gap statistics identified 6 as the optimal number of clusters (panel B, leftmost plot). Hierarchical clustering into 6 clusters yielded 5 primary clusters and 1 “noise” cluster containing only one sample (the leftmost brain). Most samples (91.8%; 101 brains) were assigned to the same groups as in Figure 2E. **(B)** Determining the optimal number of clusters for classifying brain samples into projectomic groups. Gap and silhouette indices were used to analyze the samples and identify the optimal number of clusters (*N*) for various clustering methods. The analysis was conducted on the dataset in its original dimension (280-D) for the leftmost panel and on the NMF-dimension-reduced dataset for the remaining panels. For silhouette index-based analysis^120^, the number of clusters that produced the highest silhouette value was chosen. For gap index-based analysis^116^, we used the criterion that gap(*N*) must be ≥ [gap(*N* +1) – standard error(*N* +1)] for *N* to be selected. Once the incremental gain from increasing *N* falls within the standard error—reflecting the random variability in the gap statistic—the improvement is considered negligible, and *N* is taken as optimal. We note that the optimal number of clusters consistently remained at 5 for the dimensionality-reduced dataset across different combinations of optimal cluster number criteria and clustering techniques. The error bars in the gap index plots represent the standard error of the means. GMM, Gaussian mixture model. **(C)** To evaluate the variability in clustering results using different clustering methods described in panel B, brain samples were visualized in 2-D plots. The leftmost plot (data from Figure 2E) shows results based on hierarchical clustering, and the outcomes from three other clustering methods are also represented using the same tSNE axes. As is evident, most brains consistently fall into the same group across all clustering methods. For example, the hierarchical- and *k*-means clustering methods differ by a single brain (arrow), which is assigned to the MI group by hierarchical clustering but to the BSLT group by *k*-means clustering. **(D)** Testing the sensitivity of clustering results to sampling. To evaluate the stability of the results shown in Figure 2 to specific samples in our dataset, a random 22 samples (20% of the 110 samples) were removed, and the analysis was conducted on the remaining 88 samples for each trial; this was repeated 250 times. The optimal numbers of basis patterns and clusters are summarized in the graph. Most trials (79.2%, 198 trials) yielded five projectomic groups, with 62.8% (157 trials) yielded five basis patterns and five projectomic groups.

**Figure S3.**
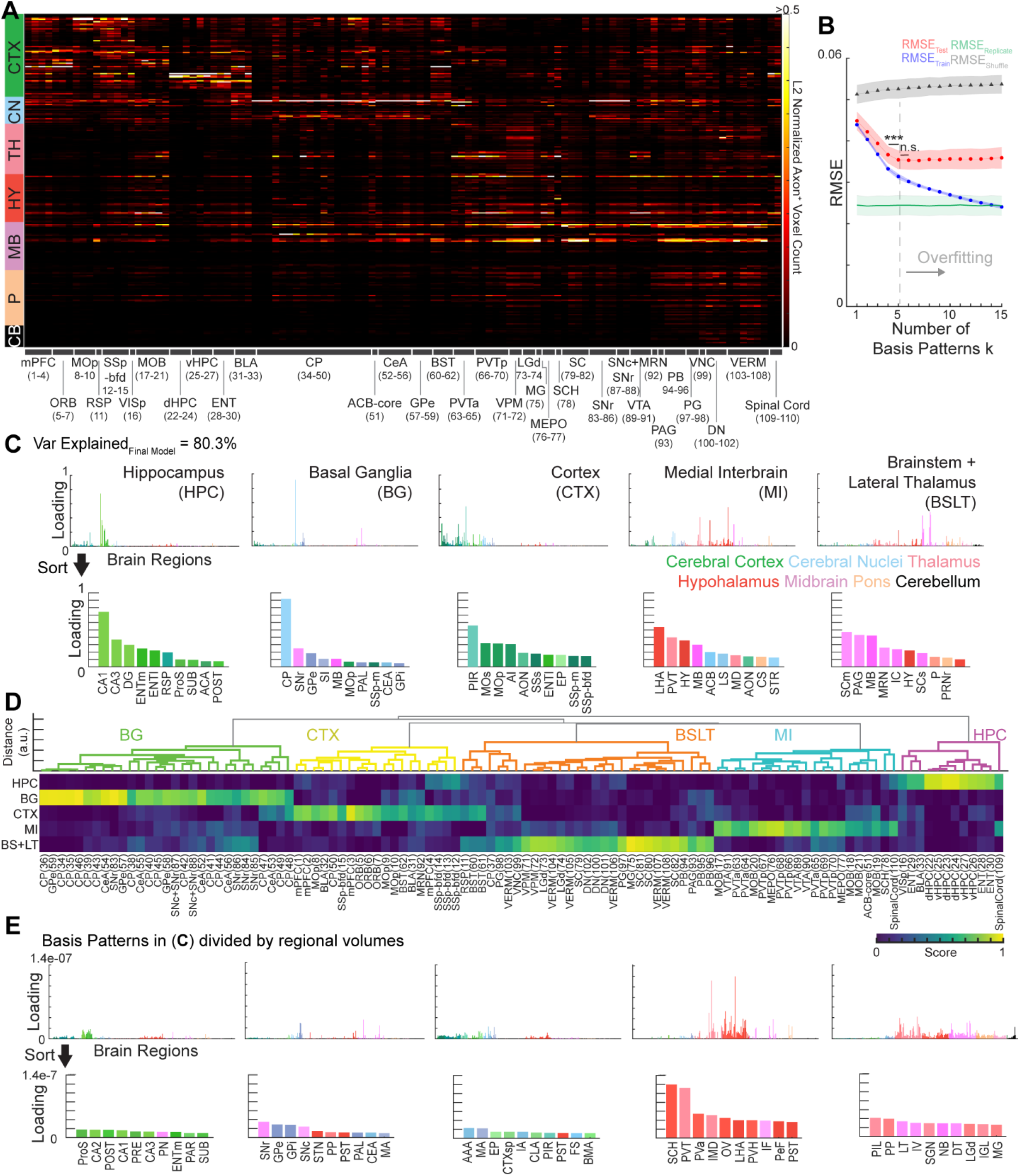
Regional axonal volumes–based NMF analysis, related to Figure 2. **(A)** Data from Figure 1D, originally plotted using normalized regional axon density, is replotted here using normalized regional axon volumes, representing the total quantity of axons in individual brain regions. **(B–D)** NMF analysis based on normalized regional axon volume data. Cross-validation results again indicated that the dataset dimension could be optimally reduced to *k* = 5 (**B**), and the projection basis patterns captured anatomical characteristics (**C**). Hierarchical clustering results (**D**) were largely consistent with those in Figure 2E, though some differences were observed. For example, SNr-injected brains were now classified within the BG group, whereas most were assigned to the BSLT group in the density-based analysis. Overall, 84.6% (93) brains were classified in the same groups as in Figure 2E. **(E)** The basis patterns shown in panel C were divided by regional volumes, yielding results similar to those obtained from the density-based analysis shown in Figure 2D.

**Figure S4.**
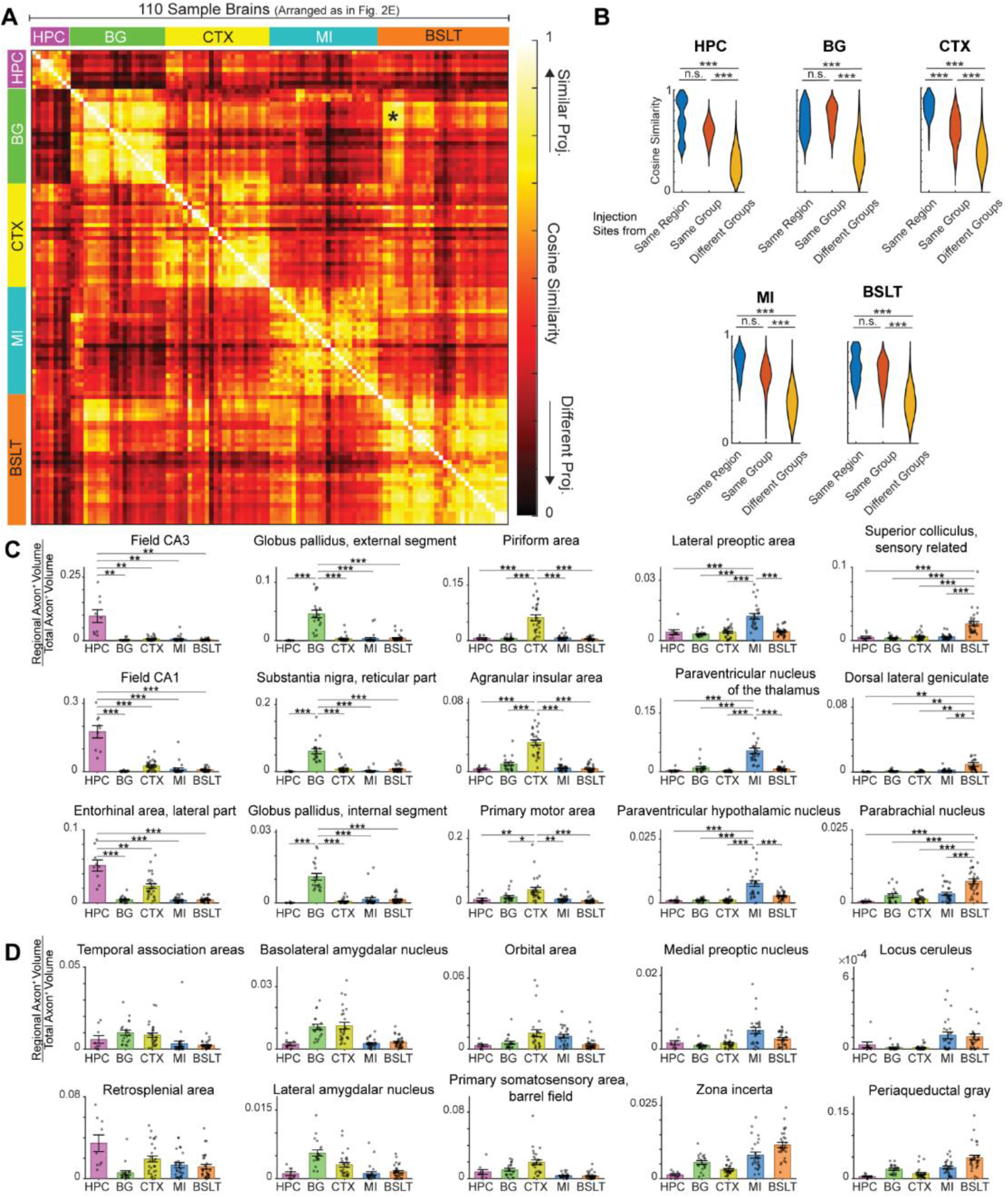
Projection similarity analysis supports NMF-based grouping and identifies marker regions, related to Figure 2. **(A)** Pairwise similarity was measured using cosine similarity on the 280-D dataset shown in Figure 1D. The brain sample order follows the clustering order presented in Figure 2E. The brains in the BSLT group forming the off-diagonal “island” (marked with *) were injected into SNr and SNc+SNr and showed high similarity to some CeA- and GPe-injected brains in the BG group. **(B)** Pairwise similarity measured between brains injected in the same regions (blue), different regions in the same projectomic groups (orange), and different regions in other projectomic groups (yellow) were statistically compared. Kruskal-Wallis test with Dunn’s post-hoc correction. ***, *p* < 0.001; n.s., not statistically significant. **(C)** Normalized axonal projection density (fraction of total axonal output per brain) across five projectomic groups (HPC, BG, CTX, MI, BSLT; n = 9, 19, 27, 25, 30 brain samples, respectively). Bars, mean ± SEM; points, individual mice. Marker regions were defined by three criteria: (1) significant group effect by one-way ANOVA (p < 0.05); (2) the highest-mean group exceeding twice the mean of every other group; and (3) Welch’s two-sided t-tests of the highest-mean group versus each of the other four groups all significant after within-region BKY FDR correction (p < 0.05). *p < 0.05, **p < 0.01, ***p < 0.001. **(D)** Examples of regions showing relatively high axonal projection density in more than one projectomic group. Normalized axonal projection density across five projectomic groups. These regions did not meet the marker-region criteria in (C); specifically, they had at least two groups whose mean exceeded half of the highest group mean.

**Figure S5.**
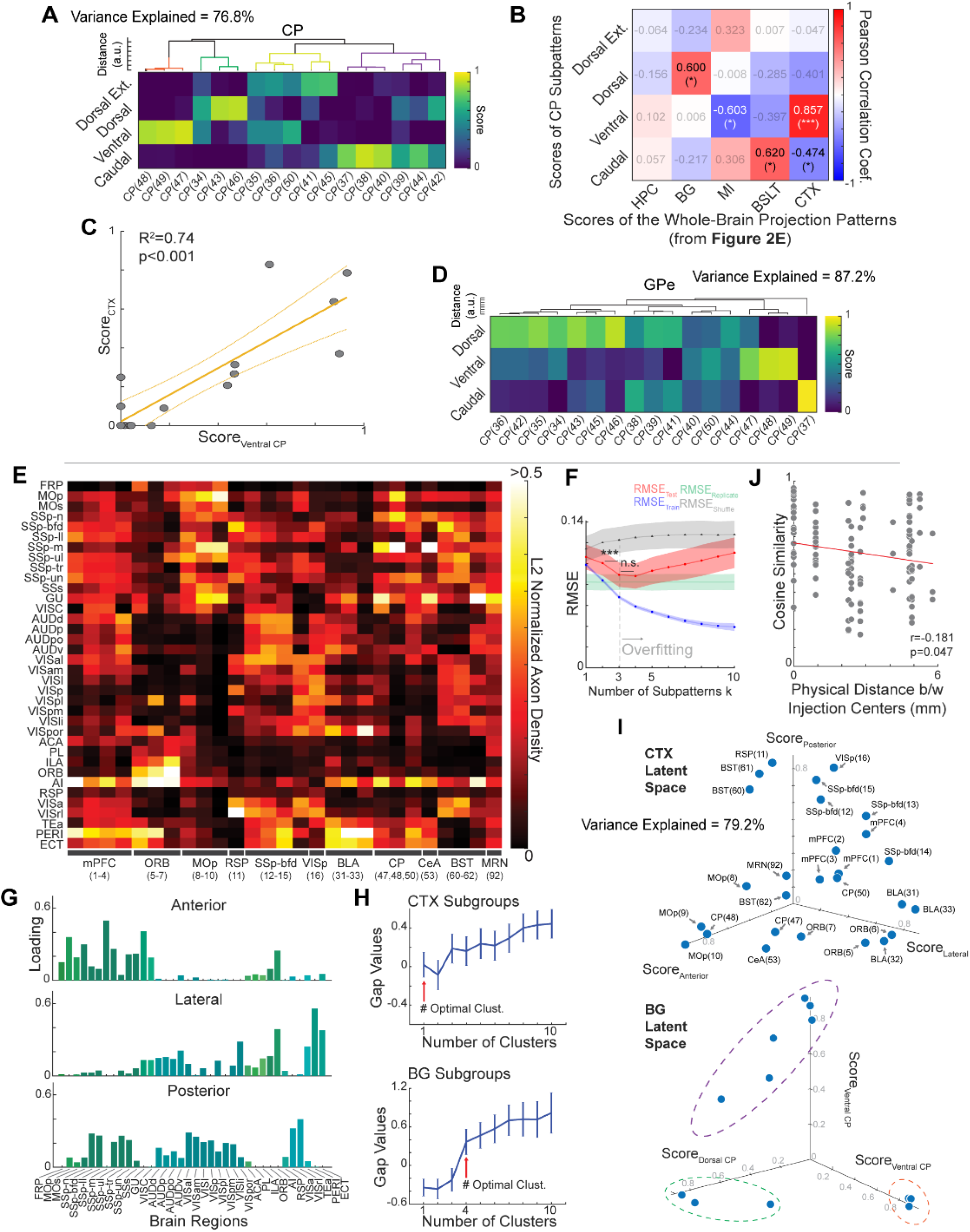
Analysis of basal ganglia and cortical subpatterns, related to Figure 3. **(A)** Axon distributions of CP-injected brains were dimensionality-reduced and hierarchically clustered using the 4D scores. Subgroups were visually distinguished by assigning unique colors to each in the dendrogram. The corresponding subgroups were enclosed within color-matching dotted circles in the bottom of Panel I. **(B)** Pairwise Pearson correlation between the scores of CP subpatterns from Panel A and scores of the whole-brain projection patterns of the 17 CP-injected brains (from Figure 2E). By extending our analysis to examine whole-brain serotonergic innervation using the scores of the CP-injected brains shown in Figure 2E, we identified statistically significant correlations between several CP subpatterns and the main projectomic patterns. *, *p* < 0.05; ***, *p* < 0.001; gray values indicate no statistical significance. **(C)** Linear regression between the ventral CP subpattern scores and the CTX pattern scores of the 17 CP-injected brains. The near-zero *y-*intercept and high R^2^ value indicates that serotonergic axon labeling in the cortex of CP-injected brains was primarily driven by the labeling of serotonergic axons targeting the ventrolateral CP. **(D)** Hierarchical clustering of GPe axon projection patterns from the brains with injections in CP. High-dimensional axon distributions were first decomposed into a low-dimensional space using the basis patterns in Figure 3E, and the resulting scores were hierarchically clustered. **(E)** L2-normalized serotonergic axon density in the isocortical regions of CTX group samples (n = 27). Injection sites are listed on the *x*-axis, and cortical projection regions are listed on the *y*-axis. **(F)** Speckled cross-validation of the axon density matrix shown in panel A indicates that the optimal number of CTX subpatterns is 3. For each iteration, 10% of randomly selected entries were used as test sets and 90% as training sets. 2500 iterations. One-way ANOVA followed by Bonferroni correction. Error bars, standard deviation. ***, *p* < 0.001; n.s., not statistically significant. **(G)** Three extracted CTX basis subpatterns showing their loadings across cortical areas. **(H)** Gap indices were applied to identify the optimal number of subgroups within the CTX (and BG groups as a comparison) using hierarchical clustering (top and bottom panels, respectively). See **Figure S2B** for details on the gap index criteria. **(I)** 3D loadings of the dimensionality-reduced individual brains in the CTX group (top) and BG group (bottom, as a comparison) are shown in their respective latent space. CTX samples are widely distributed across the latent space and do not form discrete clusters, consistent with Gap analysis suggesting a single cluster (panel H). This contrasts with the clustering observed in the BG samples, with three subgroups outlined by dashed lines colored according to hierarchical clustering in panel A. The Dorsal-Extreme subgroup is omitted for display purpose. Note that, in the CTX-sample distribution, all three basolateral amygdala (BLA)-injected brains had high scores on the lateral subpattern, like orbitofrontal cortex (ORB)-injected brains, whereas two of the three CP-injected brains had high scores on the anterior subpattern like primary motor cortex (MOp)-injected brains. **(J)** Pairwise cosine similarity and the Euclidean distance of injection target region centers exhibit a weak but significant negative linear correlation. Data are from the 16 cortex-injected samples in Panel E. Pearson correlation was used for this analysis.

**Figure S6.**
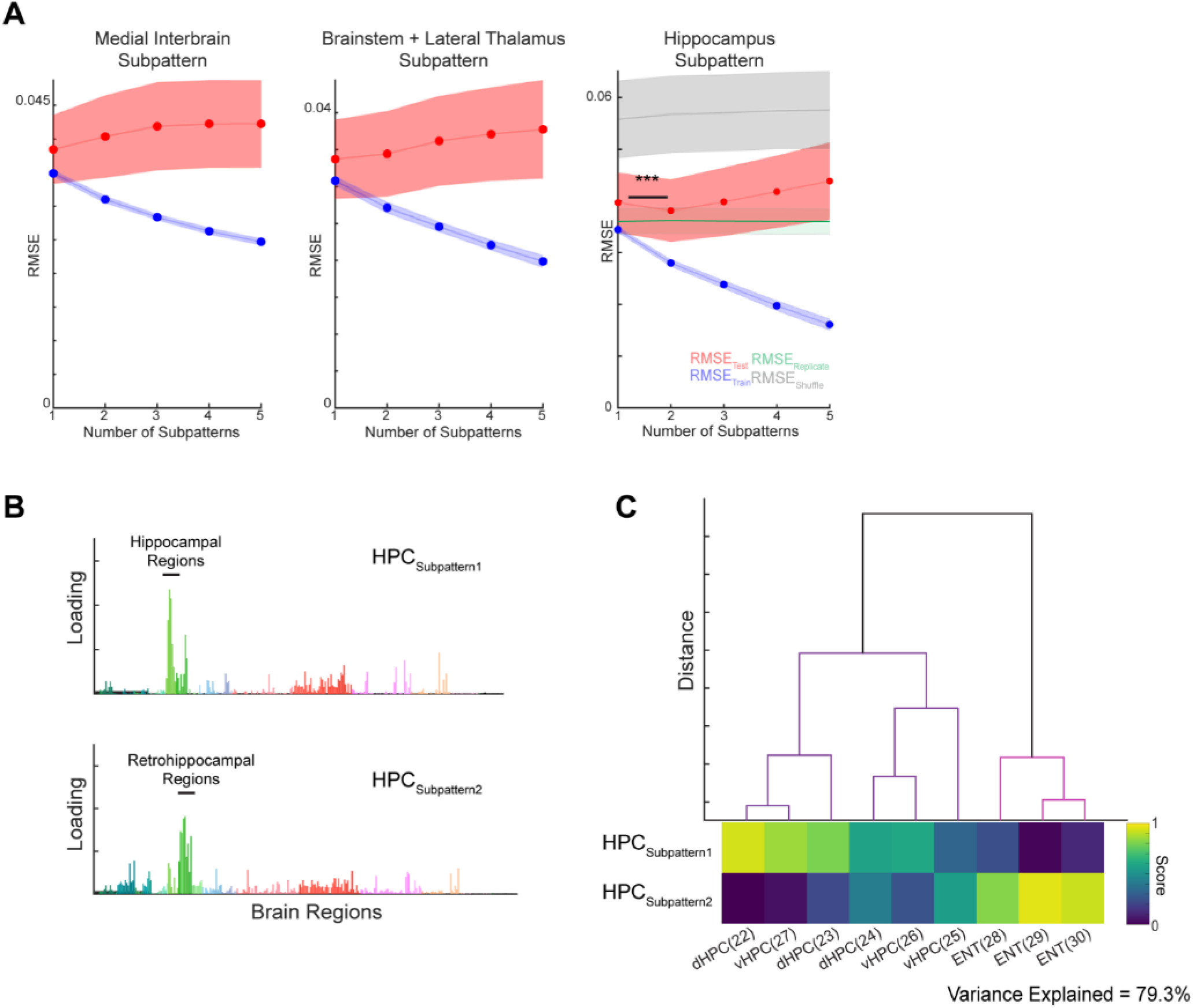
Subpattern analysis of MI, BSLT, and HPC patterns, related to Figure 3. **(A)** The same analytical pipeline from **Figure S5** was applied to the MI, BSLT, and HPC patterns. Whereas the NMF-based approach did not identify any distinct subpatterns within the MI and BSLT patterns, the HPC pattern can be divided into two subpatterns. 2500 iterations. One-way ANOVA followed by Bonferroni correction. Error bars, standard deviation. ***, *p* < 0.001. **(B)** The two HPC subpatterns exhibited high loadings in distinct regions: one in the hippocampal regions (e.g., CA1, CA2, CA3) and the other in retrohippocampal regions (e.g., entorhinal cortex). **(C)** Brains from the HPC projectomic group are hierarchically clustered based on these two identified subpatterns.

**Figure S7.**
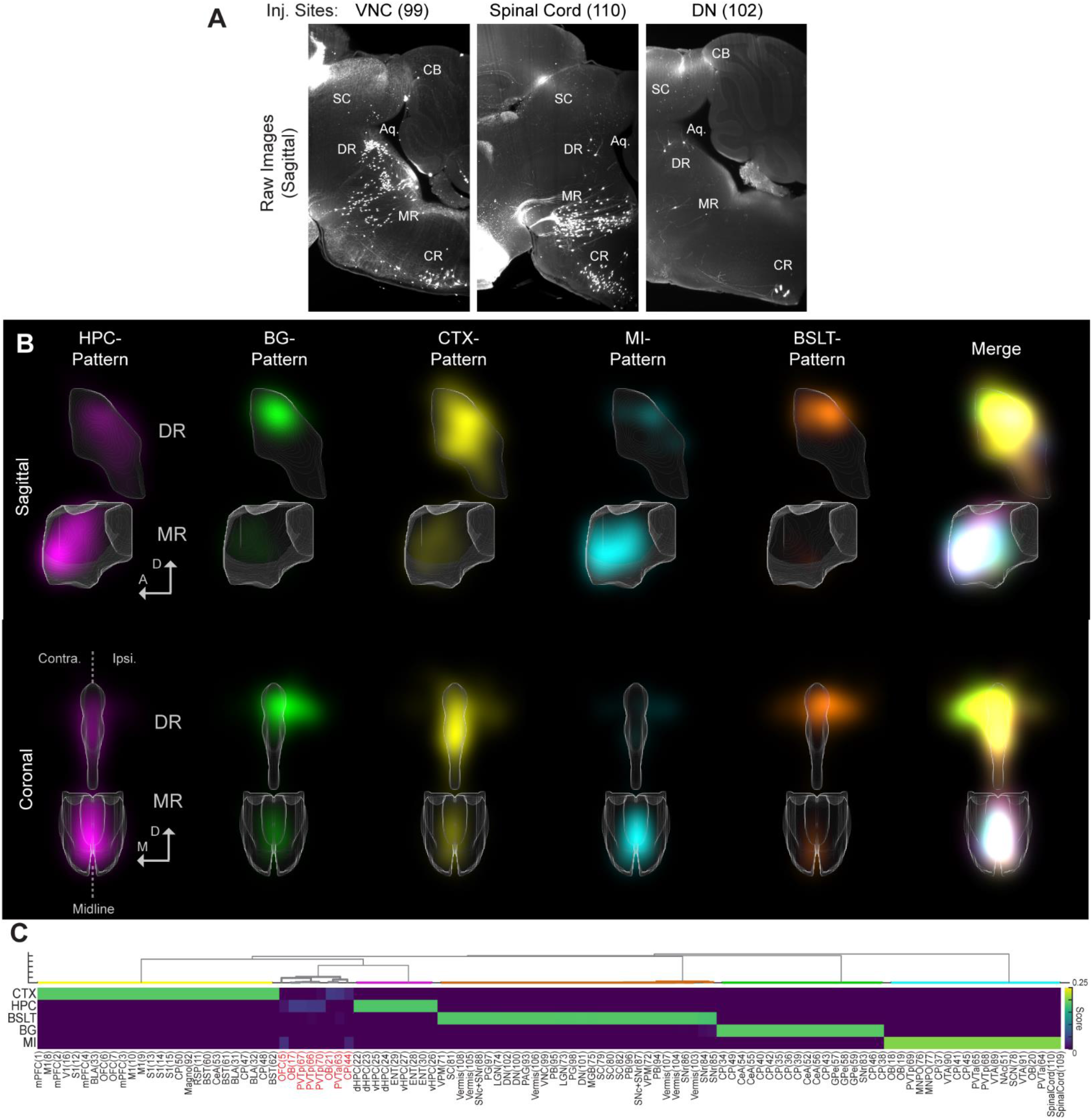
Distributions of cell body positions of serotonin neurons within DR and MR, related to Figure 4. **(A)** Example brains showing labeled serotonergic cell bodies in the caudal raphe (CR). Although we observed labeled serotonin neurons in CR in the samples injected in the hindbrain and spinal cord, they were not fully analyzed in this study, as we did not image the full distributions of these cells due to the limited imaging window size. **(B)** Supervised NMF was used to decompose cell body distributions in the DR and MR into a set of basis patterns. The optimal number of patterns was determined to be five via a hyperparameter search using Bayesian optimization with 5-fold cross-validation, a result consistent with the number of projectomic patterns. The classification accuracy reported in Figure 4C was evaluated using 5-fold stratified cross-validation with these optimized hyperparameters. For visualization, the basis patterns shown here and the sample scores in (**C**) are from a model trained on all 110 samples using the same optimized hyperparameters. **(C)** Cell body distributions in individual sample brains represented using the basis distribution patterns. Some of the brains—brain ID numbers 5, 17, 67, 66, 70, 21, 63, and 44 (labeled in red)—were not adequately described by our model.

**Figure S8.**
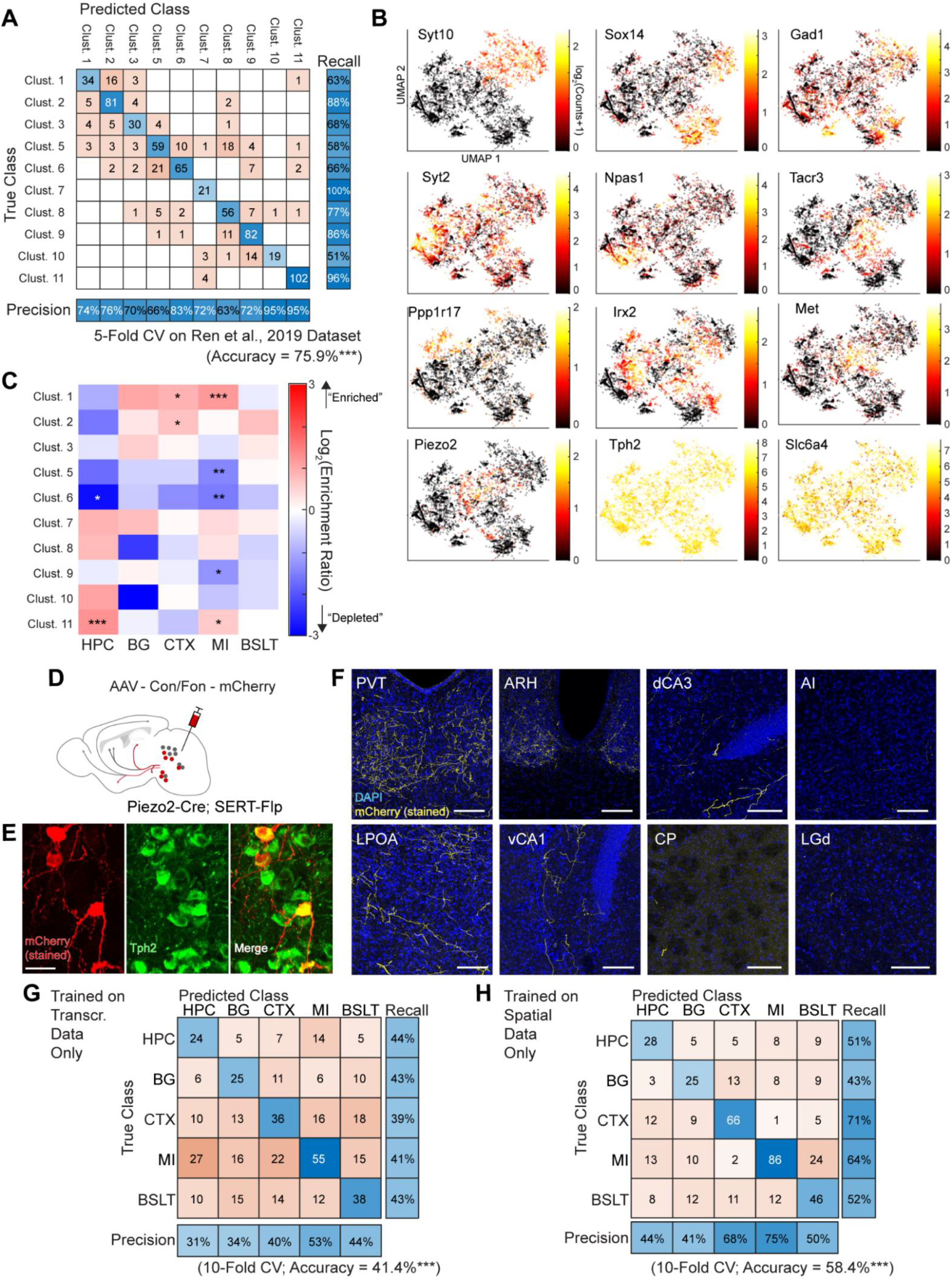
Spatial transcriptomic analysis using STARmap, related to Figure 5. **(A)** Validation of the consensus classification strategy on the published scRNA-seq dataset^25^. A classifier trained on the reference data was evaluated using 5-fold cross-validation, achieving 75.9% overall accuracy across 10 transcriptomic subtypes. Confusion matrix shows predicted versus true cluster assignments. ***p < 0.001, permutation test, 1,000 iterations (chance level = 12.3%). **(B)** Expression levels of the 10 subtype marker genes across the 10 transcriptomic clusters identified by STARmap, confirming consistency with expected expression patterns from the reference dataset. **(C)** Enrichment ratio (ER) heatmap showing the association between transcriptomic clusters and projectomic groups. Color indicates ER, defined as the proportion of a transcriptomic cluster among a group’s barcode-positive cells divided by its proportion in the overall population. Red indicates enrichment (ER > 1); blue indicates depletion (ER < 1). Asterisks denote statistical significance (Fisher’s exact test, FDR-corrected; *p < 0.05, **p < 0.01, ***p < 0.001). **(D)** Intersectional validation strategy. A Cre/Flp-dependent mCherry reporter (AAV-Con/Fon-mCherry) was injected into the DR and MR of *Piezo2-Cre;SERT-Flp* mice to selectively label Piezo2-expressing serotonin neurons. **(E)** mCherry-labeled neurons in the raphe co-express *Tph2*, confirming their serotonergic identity. Of 102 mCherry⁺ neurons identified across 3 mice (2 females, 1 male), 90 (88.2%) were *Tph2*-positive. Scale bar, 25 μm. **(F)** Projection targets of *Piezo2*⁺ serotonin neurons. mCherry-positive axons were observed in the MI- (e.g., PVT, ARH, LPO) and hippocampal regions (e.g., dCA3, vCA1) but nearly absent in the projection targets of CTX, BG, and BSLT groups (e.g. AI, CP, LGd, respectively) in all 3 mice examined, consistent with the enrichment of Cluster 11 (*Met*/*Piezo2*-high) neurons in the HPC- and MI-projecting groups. Scale bars, 100 μm. (**G**, **H**) Confusion matrices from 10-fold cross-validated random forest classifiers trained on transcriptomic features only (10 marker genes, *Tph2, Sert*; panel G) or spatial features only (AP, ML, DV positions, and DR/MR region; panel H). Class weighting was applied for unequal group sizes. ***p < 0.001, permutation test, 1,000 iterations (chance level = 20.0% for transcriptomic, 21.1% for spatial). Note that the cell-level spatial accuracy (58.4%) is lower than the animal-level distribution-based prediction (83.6%; Figure 4C), reflecting that single-cell spatial coordinates carry less information about projectomic identity than the full cell body distribution within an animal.

**Figure S9.**
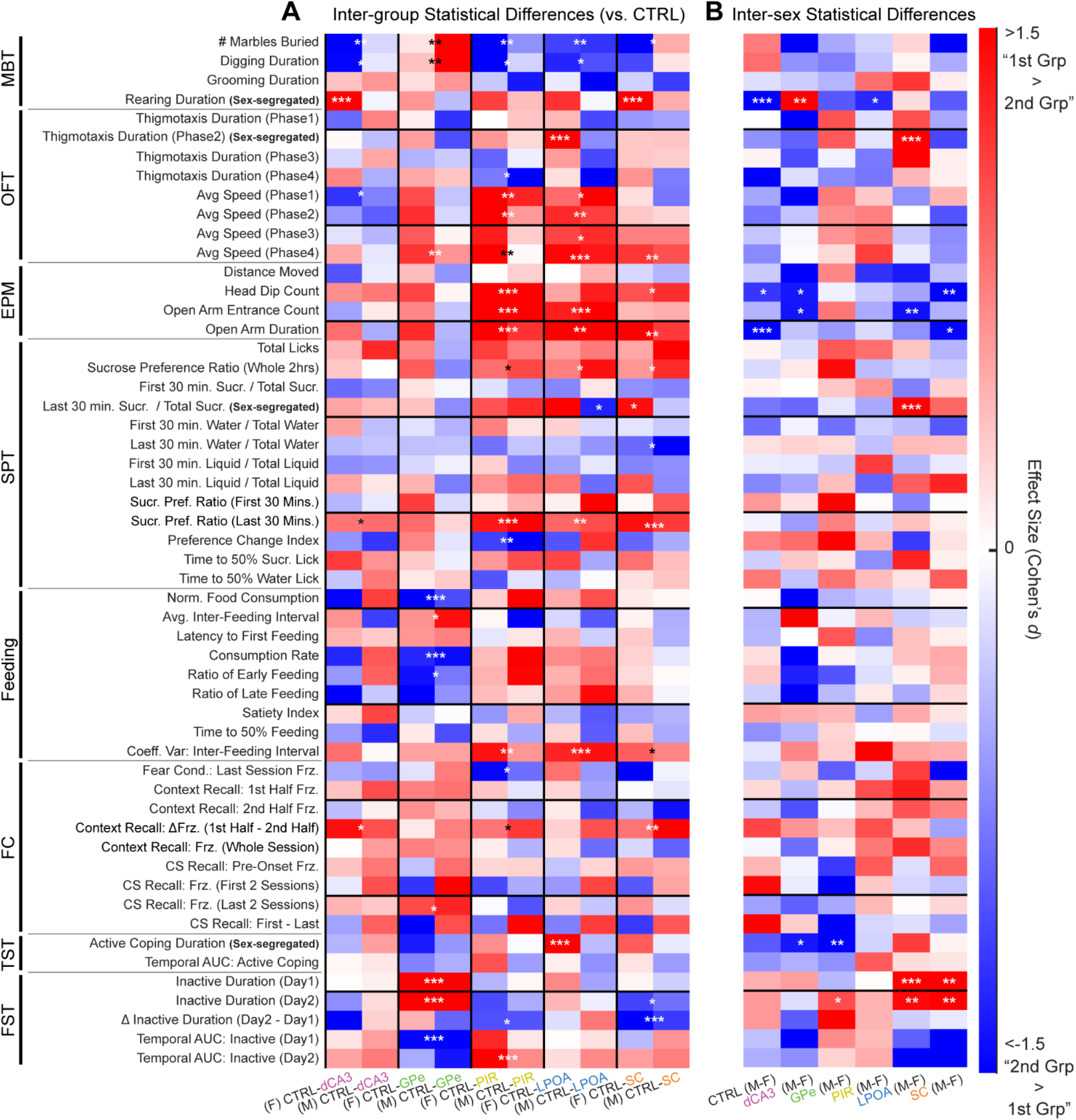
Statistical comparisons and effect sizes across 54 behavioral features, related to Figure 6. **(A)** Pairwise post-hoc tests against the control group were performed conditionally based on ANOVA results. See **Methods** for detailed descriptions of the 54 behavioral features. Inter-group comparisons were conducted only if a significant main effect of group was present (*p*_group_ < 0.05). If the interaction (sex × group) effect was significant (*p*_interaction_ < 0.05), analyses were stratified by sex (comparing experimental males with control males, and females with control females). If the interaction was not significant, sexes were pooled within groups. Regardless of statistical significance, Cohen’s *d* effect sizes 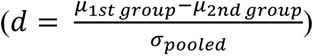 are represented by a red-white-blue color scale to show the magnitude and direction of differences in a sex-specific manner. For example, the blue shading of "# Marbles Buried" in the "(F) CTRL-dCA3" column denotes that the female dCA3-injected group buried more marbles than the female control group. BKY FDR-corrected across the 5 group-vs-control comparisons within each behavioral feature. \**p*_corrected_<0.05; \*\**p*_corrected_<0.01; \*\*\**p*_corrected_<0.001. **(B)** Likewise, within-group sex differences were statistically tested only if the main effect of sex or the interaction effect was significant (*p*_sex_ or *p*_interaction_ < 0.05).

**Figure S10.**
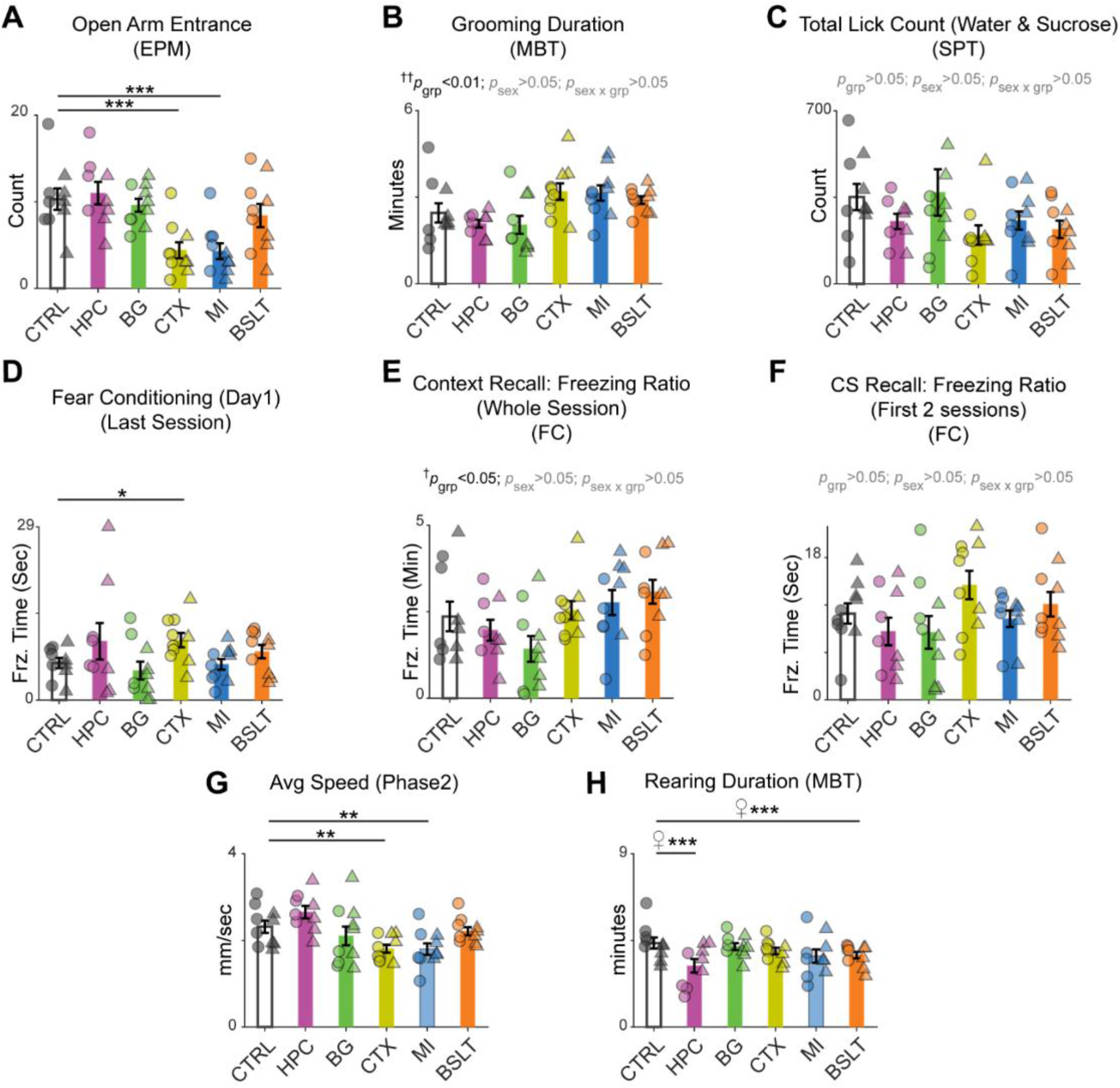
Extended behavioral characterization and feature correlation structure, related to Figure 6. **(A–H)** Additional representative behavioral features. Note that in Panel B, despite a significant group effect detected by two-way ANOVA, no pairwise differences relative to control were identified in the post hoc comparisons. In Panel H, sex-stratified pairwise comparisons were conducted as post hoc tests, as the interaction p-value was significant (*p*_sex-group_ < 0.01). \**p*_corrected_ < 0.05; \*\**p*_corrected_ < 0.01; \*\*\**p*_corrected_ < 0.001; ^†^*p*_group_ < 0.05; ^††^*p*_group_ < 0.01

**Figure S11.**
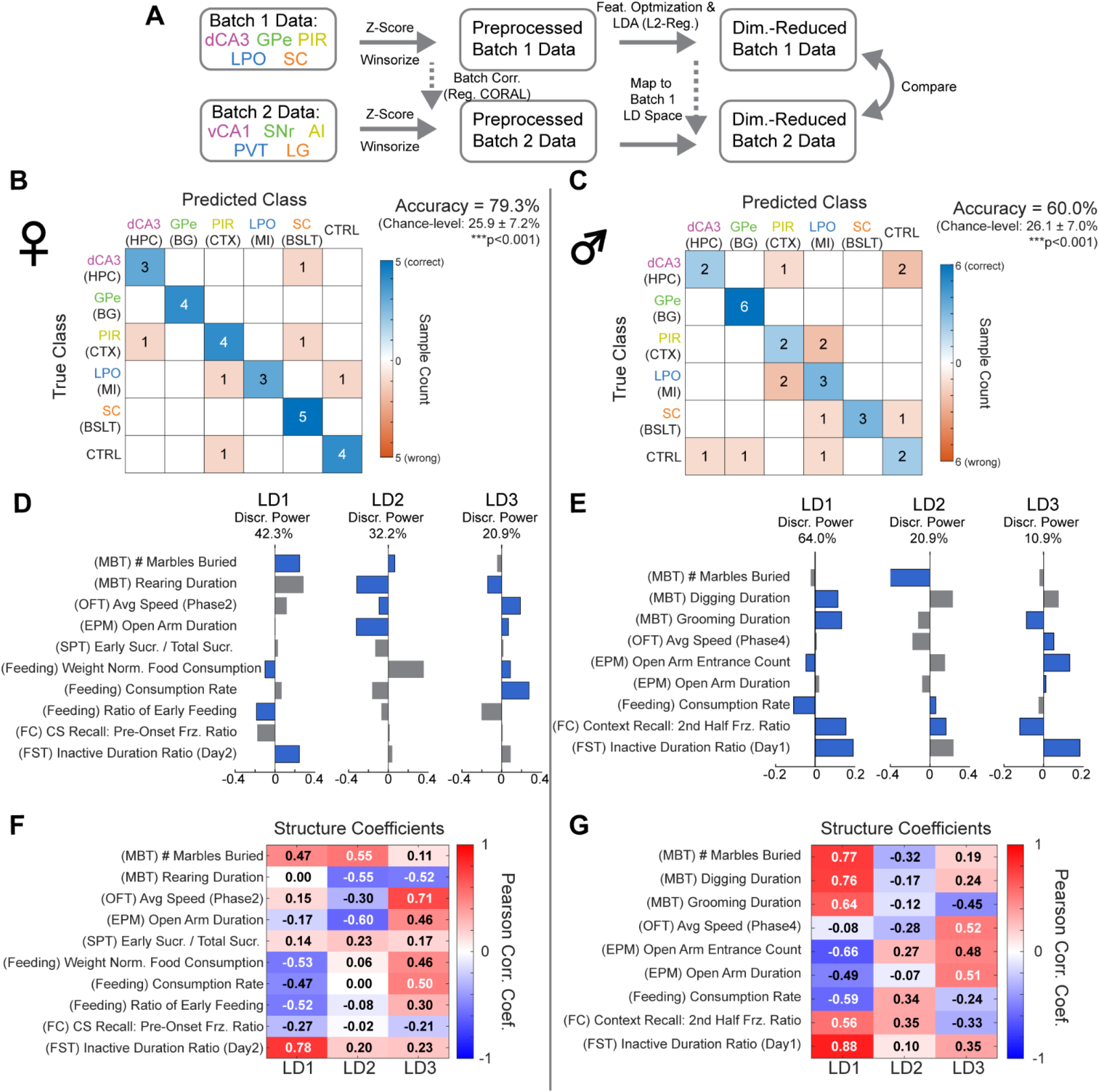
Supervised dimensionality reduction pipeline, model performance, and LD axis interpretation, related to Figure 7. **(A)** Multidimensional model-based analysis pipeline. Dimensionality reduction was performed on Batch 1 data using L2-regularized LDA (top row). Using parameters derived from the Batch 1 model, Batch 2 samples were projected onto the Batch 1 LD space. See **Methods** for details. **(B, C)** Performance of the model used for downstream analyses. A nearest-centroid classifier was applied to Batch 1 LD scores obtained using the selected feature sets and LD components. Validation was performed using leave-one-out cross-validation (LOOCV). (**D**, **E**) A hyperparameter search to select optimal feature sets and the number of LD components identified three LDs for both sexes. Displayed are the features selected to effectively segregate groups, along with feature loadings derived from all Batch 1 samples using the optimized hyperparameters. Blue bars indicate features with an absolute structure coefficient (shown in Panels F and G) greater than 0.3, suggesting a substantive association with the discriminant axis. (**F, G**) Structure coefficients calculated as Pearson correlations between individual behavioral features and the LD scores. These coefficients quantify the direct association between each behavioral feature and LDs— independent of inter-feature correlations—and identify the primary behavioral drivers of group separation together with the LD loadings shown in Panels D and E.

**Figure S12.**
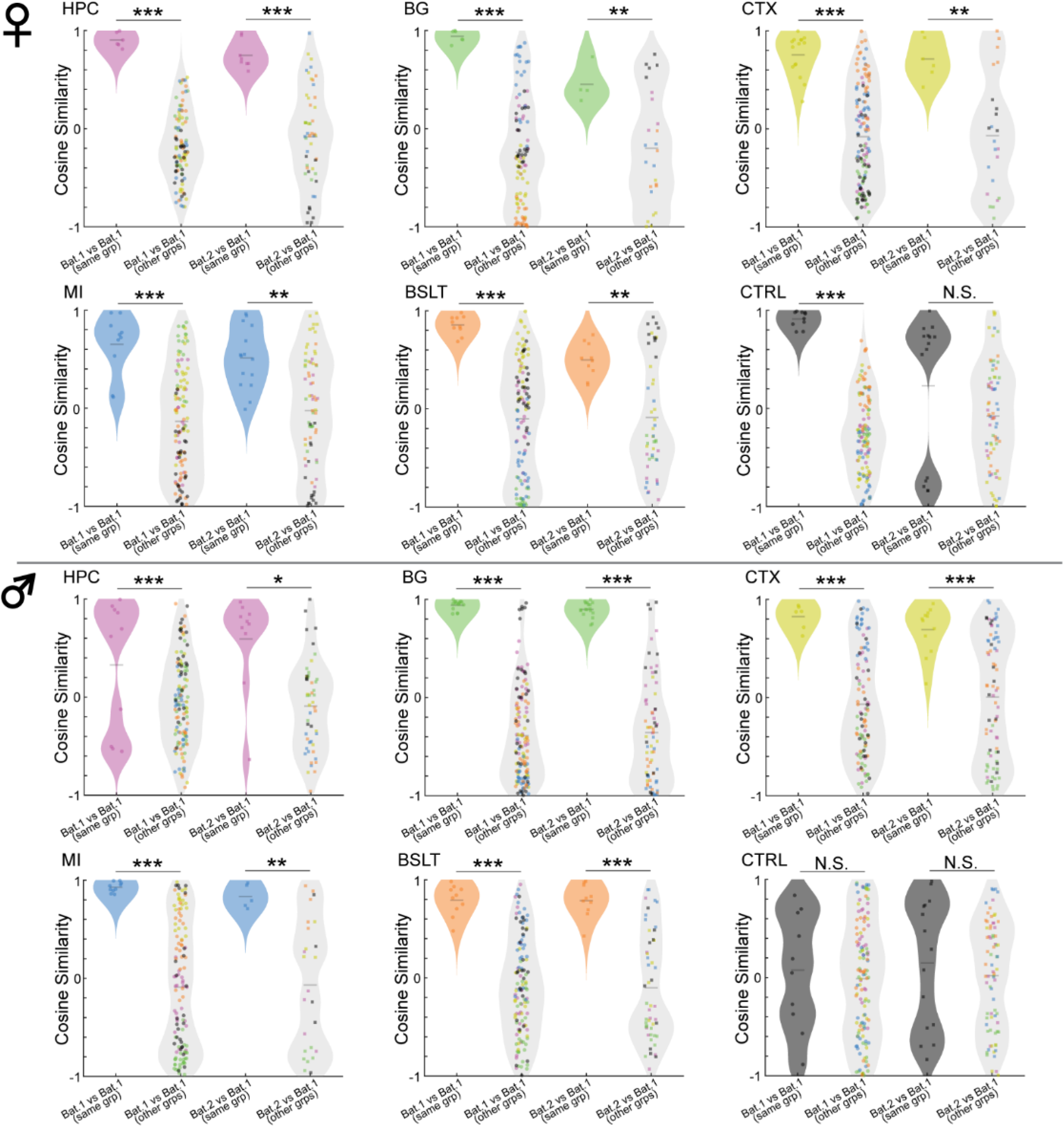
Within- and between-group cosine similarity of behavioral phenotypes across batches, related to Figure 7. Cosine similarity was computed from the linear discriminant (LD) scores shown in Figure 7G and H, to quantify how consistently serotonergic projection-based groups cluster by behavioral phenotype within and across batches. Within-group similarity (colored violins) reflects pairs drawn from the same projection group, either within the same batch or across batches; between-group similarity (gray violins) reflects pairs drawn from the group of interest and samples outside that group. Higher within-group relative to between-group similarity indicates that serotonergic projectomic groupings generalize across batches. Each dot represents a single pairwise cosine similarity value and is color-coded according to the projectomic group identity of the comparison sample (e.g., green dots indicate pairs involving BG-group samples). Within- vs. between-group cosine similarities were compared using a two-sided permutation test on the difference of means (n = 10,000 permutations). Multiple comparisons were corrected using the Benjamini-Hochberg FDR procedure. \**p* < 0.05, \*\**p* < 0.01, \*\*\**p* < 0.001.

